# TRIM37 prevents formation of centriolar protein assemblies by regulating Centrobin stability

**DOI:** 10.1101/2020.09.03.281097

**Authors:** Fernando R. Balestra, Benita Wolf, Andrés Domínguez-Calvo, Alizée Buff, Tessa Averink, Marita Lipsanen-Nyman, Coralie Busso, Pablo Huertas, Rosa M. Ríos, Pierre Gönczy

## Abstract

TRIM37 is an E3 ubiquitin ligase mutated in Mulibrey nanism, a disease characterized by impaired growth and increased tumorigenesis, whose cellular etiology is poorly understood. TRIM37 depletion from tissue culture cells results in supernumerary foci bearing the centriolar protein Centrin. Here, we characterized these centriolar protein assemblies (Cenpas) to uncover the mechanism of action of TRIM37. We established that an atypical *de novo* assembly pathway is notably involved in forming Cenpas, which can nevertheless trigger further centriole assembly and act as MTOCs. We found also that Cenpas are present and act similarly in Mulibrey patient cells. Through correlative light electron microscopy, we uncovered that Cenpas correspond to centriole related structures and elongated electron-dense structures with stripes. Importantly, we established that TRIM37 regulates the stability and solubility of the centriolar protein Centrobin. Our findings suggest that elongated Centrobin assemblies are a major constituent of the striped electron dense structures. Furthermore, we established that Cenpas formation upon TRIM37 depletion requires PLK4 activity, as well as two parallel pathways relying respectively on Centrobin and PLK1. Overall, our work uncovers how TRIM37 prevents the formation of Cenpas that would otherwise threaten genome integrity, including possibly in Mulibrey patients.

## INTRODUCTION

Centrioles are tiny evolutionarily conserved cylindrical organelles characterized by nine triplets of microtubules (MTs) arranged with a striking 9-fold radial symmetry (reviewed in (Gönczy, 2012; Gönczy and Hatzopoulos, 2019)). In addition to MTs, centrioles contain multiple copies of tens of distinct proteins that contribute to their assembly, structure and function (Andersen et al., 2003; Jakobsen et al., 2011). Centrioles are essential for the formation of cilia and also recruit pericentriolar material (PCM), including the MT nucleator γ-tubulin ring complex, thus forming the centrosome of animal cells (reviewed in (Bornens, 2012)). Probably because of such important roles, centriole number is tightly regulated, with most cycling cells having two units at the cell cycle onset and four units by the time of mitosis (reviewed in (Sullenberger et al., 2020)). Alterations in centriole number control can have an adverse impact on cell physiology and genome integrity. Thus, supernumerary centrioles lead to extra cilia and centrosomes (Duensing et al., 2007; Habedanck et al., 2005; Mahjoub and Stearns, 2012), which can be observed also in several human disease conditions, including certain cancer types (reviewed in (Bettencourt-Dias et al., 2011; Chavali et al., 2014; Gönczy, 2015; Nigg and Holland, 2018; Nigg and Raff, 2009). Despite their importance, the mechanisms that prevent the formation of excess centriolar structures remain incompletely understood.

The two centrioles present at the onset of the cell cycle differ in age: whereas the older, mother, centriole is at least two cell generations old, the younger, daughter, centriole was formed in the previous cell cycle. The mother centriole bears distinctive distal and sub-distal appendages that the daughter centriole acquires only later during the cell cycle (reviewed in (Sullenberger et al., 2020)). In human cells, the proximal region of both mother and daughter centrioles in the G1 phase of the cell cycle is encircled by a torus bearing the interacting proteins CEP57/CEP63/CEP152 (Brown et al., 2013; Lukinavicius et al., 2013) (reviewed in (Banterle and Gönczy, 2017)). The Polo-like-kinase PLK4 is recruited to this torus, where it focuses to a single location towards the G1/S transition, owing notably to a protective interaction with its substrate STIL, thus marking the site of procentriole assembly (Klebba et al., 2015; Moyer et al., 2015; Ohta et al., 2014) (reviewed in (Arquint and Nigg, 2016)). The onset of procentriole assembly entails formation of a 9-fold radially symmetric cartwheel thought to act as a scaffold for the organelle (reviewed in (Guichard et al., 2018; Hirono, 2014)). The fundamental building block of the cartwheel is HsSAS-6, which self-assembles *in vitro* into structures akin to those found *in vivo* (Guichard et al., 2017; Kitagawa et al., 2011b; Strnad et al., 2007; van Breugel et al., 2011). During S/G2, the emerging procentriole remains closely associated with the resident centriole and elongates through the contribution notably of the centriolar proteins CPAP/SAS-4, SPICE as well as C2CD3 (Balestra et al., 2013; Comartin et al., 2013; Kohlmaier et al., 2009; Schmidt et al., 2009; Tang et al., 2009; Thauvin-Robinet et al., 2014). During mitosis, the procentriole disengages from the resident centriole in a manner that requires the activity of the Polo-like-kinase PLK1, with increased PLK1 levels during S/G2 leading to premature centriole disengagement and centriole reduplication (Loncarek et al., 2010; Tsou et al., 2009). Normally, disengagement during mitosis generates two centriolar units that are then licensed to recruit PCM and trigger a new round of centriole assembly in the following cell cycle.

Centrioles can also assemble independently of a resident centriole. Such *de novo* assembly can occur in physiological conditions, for instance when the protist *Naegleria gruberii* transitions from an acentriolar amoeboid life form to a flagellated mode of locomotion (Fritz-Laylin et al., 2016; Fulton and Dingle, 1971). Likewise, centrioles assemble *de novo* at the blastocyst stage in rodent embryos (Courtois et al., 2012). *De novo* assembly of centrioles can also be triggered experimentally in human cells following removal of resident centrioles through laser ablation or chronic treatment with the PLK4 inhibitor Centrinone (Khodjakov et al., 2002; Wong et al., 2015). Therefore, in human cells, *de novo* assembly is normally silenced by resident centrioles. In contrast to the situation in physiological conditions, experimentally provoked *de novo* centriole assembly in human cells is error prone and lacks number control (La Terra et al., 2005; Wong et al., 2015). Moreover, *de novo* assembly of foci that contain some centriolar proteins and which can function as MTOCs forms in human cells upon depletion of the intrinsically disordered protein RMB14 or the Neuralized Homology repeat containing protein Neurl4 (Li et al., 2012; Shiratsuchi et al., 2015). Such extra foci, although not *bona fide* centrioles as judged by electron-microscopy, threaten cell physiology and could conceivably contribute to disease.

TRIM37 is a RING-B-box-coiled-coil protein with E3 ubiquitin ligase activity (Kallijarvi et al., 2002; Kallijarvi et al., 2005), which somehow prevents the formation of foci bearing centriolar markers (Balestra et al., 2013). Individuals with loss of function mutations in both alleles of TRIM37 are born with a rare disorder known as Mulibrey nanism (Muscle-liver-brain-eye nanism). The main features of this disorder are growth failure with prenatal onset, as well as characteristic dysmorphic features and impairment in the organs that give rise to the name of the condition (Avela et al., 2000). In addition, Mulibrey patients have a high probability of developing certain tumor types (Karlberg et al., 2009). Mice lacking Trim37 recapitulate several features of Mulibrey nanism, including a higher propensity to form tumors (Kettunen et al., 2016). However, the cellular etiology of Mulibrey nanism remains unclear, partially because of the many roles assigned to this E3 ubiquitin ligase. In tissue culture cells, TRIM37 mono-ubiquitnates and thereby stabilizes PEX5, promoting peroxisomal function (Wang et al., 2017). However, Trim37 knock out mice and mouse cell lines depleted of Trim37 do not exhibit peroxisomal associated phenotype (Wang et al., 2017), suggesting that the conserved pathological features exhibited also by the mouse disease model must have a different cellular etiology. Furthermore, the chromosomal region 17q23 where TRIM37 resides is amplified in ~40% of breast cancers (Sinclair et al., 2003). TRIM37 mono-ubiquitinates histone H2A in the MCF-7 breast cancer cell line dampening the expression of thousands of genes, including tumors suppressors, thus offering a potential link between TRIM37 overexpression and tumorigenesis (Bhatnagar et al., 2014). Furthermore, TRIM37 overexpression has been linked to increased cell invasion and metastasis in colorectal and hepatocellular carcinoma (Hu and Gan, 2017; Jiang et al., 2015). Therefore, both the depletion and the excess of TRIM37 are accompanied by detrimental consequences.

We previously performed a genome wide siRNA-based screen in human cells to identify regulators of centriole assembly, using the number of foci harboring the centriolar marker Centrin-1:GFP as a readout. In this screen, we identified TRIM37 as a potent negative regulator of Centrin-1:GFP foci number. Our initial characterization of the TRIM37 depletion phenotype revealed that ~50% of cells possess supernumerary foci harboring the centriolar proteins Centrin and CP110, as well as instances of multipolar spindle assembly and chromosome miss-segregation. Additionally, we found that inhibition of PLK1 partially suppressed supernumerary foci formation upon TRIM37 depletion, leading to the suggestion that such foci occur through centriole reduplication (Balestra et al., 2013), although the fact that suppression was partial suggested that an additional explanation was to be found. Here, we set out to further explore the nature of such supernumerary foci to uncover the mechanism of action of TRIM37, and perhaps thereby also provide novel insights into Mulibrey nanism.

## RESULTS

### TRIM37 prevents formation of centriolar protein assemblies (Cenpas)

To further decipher the origin of supernumerary foci containing Centrin and CP110 following TRIM37 depletion, we investigated where in the cell they first appeared. We reasoned that appearance of supernumerary foci close to resident centrioles could indicate centriole reduplication, whereby premature disengagement would license resident centrioles and procentrioles to prematurely seed centriole assembly. By contrast, appearance of supernumerary foci away from resident centrioles would suggest some type of *de novo* process. We performed live imaging of HeLa cells expressing Centrin-1:GFP (referred to as HC1 cells hereafter) and depleted of TRIM37 by siRNAs. As shown in Figure 1A, we found that extra Centrin-1:GFP foci can appear in the vicinity of resident centrioles (yellow arrows, 8/13 foci), but also far from them (orange arrows, 5/13 foci). These results suggest that extra Centrin-1:GFP foci upon TRIM37 depletion may form both through centriole reduplication and some type of d*e novo* process.

**Figure 1.**
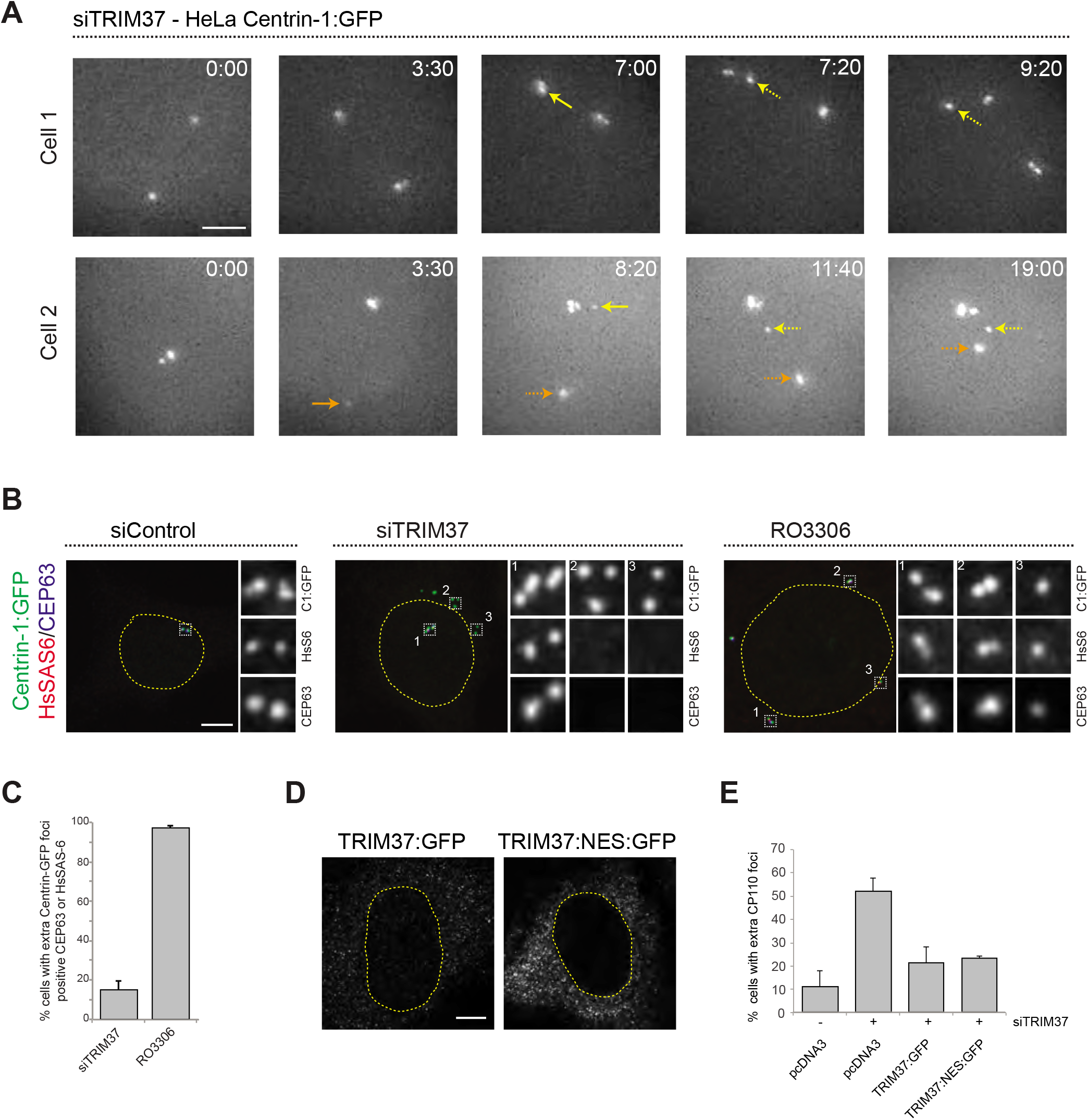
Supernumerary centriolar protein assemblies (Cenpas) form upon TRIM37 depletion. **A.** Relevant images from wide-field time-lapse recordings of HeLa cells expressing Centrin-1:GFP and depleted of TRIM37 for 48h before imaging onset (10 min. time frame). Yellow arrows point to two foci appearing close to resident centrioles (8/13 extra foci in 11 cells), orange arrow to one focus appearing away from resident centrioles (5/13 extra foci). Solid arrows indicate first occurrence of foci, dashed arrows their continued presence. Time is indicated in h:min since imaging onset. Note that the intensity of extra Centrin-1:GFP foci is typically weaker than that of regular centrioles, especially in the early assembly stages. Note also resident centriole and procentriole appearing in the field of view at the bottom right in Cell 1, 9:20. In this and other Figure panels, scale bars correspond to 5 μm, unless indicated otherwise. **B.** HeLa cells expressing Centrin-1:GFP upon treatment with control or TRIM37 siRNAs, or upon RO3306 addition for 48h. Cells were immunostained for GFP, HsSAS-6 and CEP63. Nuclear contours are drawn with dashed yellow lines. In this and subsequent figures, magnified images from indicated regions are shown. **C.** Corresponding percentage of cells with extra Centrin-1:GFP foci that also harbor CEP63 and/or HsSAS-6. Unless otherwise indicated, all graphs report averages from two or more independent experiments (n = 50 cells each), along with SDs; P<0.01 here. Note that extra Centrin-1:GFP foci could be positive for both Cep63 and HsSAS-6 in RO3306 treated cells. **D.** HeLa cells expressing TRIM37:GFP or TRIM37 tagged with a nuclear export signal and GFP (TRIM37:NES:GFP) immunostained for GFP. **E.** Quantification of extra number of CP110 foci in HeLa cells treated with control or TRIM37 siRNAs and transfected with indicated plasmids (pcDNA3: parental vector). Cells were immunostained for GFP and CP110. The difference between TRIM37:GFP and. TRIM37:NES:GFP is not significant; P=0.7327.

To further investigate this question, we analyzed fixed S/G2 HC1 cells with antibodies against GFP to monitor Centrin-1:GFP foci, as well as against CEP63 to mark the proximal region of resident centrioles and HsSAS-6 to mark procentrioles. As expected, we found that control cells harbored four Centrin-1:GFP foci, two of which were CEP63 positive and two of which were HsSAS-6 positive (Fig. 1B). Strikingly, in cells depleted of TRIM37, we found that in addition to the normal four Centrin-1:GFP foci accompanied by two Cep63 foci and two HsSAS-6 foci, ~90% of extra Centrin-1:GFP foci did not harbor CEP63 or HsSAS-6 (Fig. 1B, 1C). For comparison, we likewise analyzed cells arrested in G2 following treatment with the CDK1 inhibitor RO3306, which induces PLK1-dependent centriole reduplication (Loncarek et al., 2010). In this case, >90% of extra Centrin-1:GFP foci harbored CEP63 and/or HsSAS-6 (Fig. 1B, 1C), in contrast to the situation upon TRIM37 depletion, further indicating that TRIM37 does not act solely to prevent centriole reduplication.

Overall, we conclude that TRIM37 depletion results in extra Centrin-1:GFP foci both near and far from resident centrioles, suggestive of centriole reduplication happening together with some d*e novo* process. Moreover, we find that such foci harbor some centriolar proteins but usually not others. We will hence refer hereafter to these entities as <u>Cen</u>triolar <u>p</u>rotein <u>as</u>semblies, or Cenpas in short.

### TRIM37 regulates Cenpas formation from outside the nucleus and localizes to centrosomes

TRIM37 can regulate transcription through nuclear association with the polycomb repressive complex 2 (PRC2) (Bhatnagar et al., 2014). To explore whether TRIM37 may function as a transcriptional regulator in preventing Cenpas formation, we addressed whether rescue of the TRIM37 depletion phenotype depended on the presence of the protein in the nucleus. We generated a version of TRIM37 forced to exit the nucleus via fusion to a nuclear export signal (NES), finding that both TRIM37:GFP and TRIM37:NES:GFP equally rescued the TRIM37 depletion phenotype (Fig. 1D, 1E). This indicates that TRIM37 acts outside the nucleus to prevent Cenpas formation.

Given the TRIM37 depletion phenotype, we explored whether the protein localizes to centrioles. Since antibodies did not prove suitable to address this question (Balestra et al., 2013; Meitinger et al., 2016), we instead expressed TRIM37:GFP, finding it to be present weakly in the nucleus and more so in the cytoplasm (Fig. S1A, S1B). Intriguingly, in some cells, TRIM37:GFP also localized to centrosomes marked by γ-tubulin (Fig. S1A, S1B). To investigate whether this might reflect a cell cycle restricted distribution, TRIM37:GFP expressing cells were probed with antibodies against GFP and Centrobin, which localizes preferentially to the resident daughter centriole and to procentrioles (Zou et al., 2005). Therefore, G1 cells bear a single Centrobin focus while S/G2 cells bear 2 or 3 (Fig. S1C), which enabled us to establish that whereas only ~10% of G1 cells harbored centrosomal TRIM37:GFP, ~60% of S/G2 cells did so (Fig. S1D). We also localized the fusion protein with respect to CEP63, Centrin-2 and the distal appendage protein CEP164, finding that TRIM37:GFP partially overlapped with CEP164 (Fig. S1E, S1F). Overall, we conclude that TRIM37 localizes to the distal part of centrioles, and it will be interesting to investigate in the future whether TRIM37 acts from this location to prevent the formation of at least some Cenpas, perhaps those in the vicinity of resident centrioles.

### Cenpas can act as MTOCs, are present in Mulibrey patient cells and trigger new rounds of centriolar assembly

TRIM37 depleted cells exhibit an increased incidence of multipolar spindles and chromosome miss-segregation (Balestra et al., 2013), suggesting that Cenpas can nucleate microtubules and serve as extra microtubule organizing centers (MTOCs). To thoroughly test this possibility, we performed microtubule depolymerization-regrowth experiments. We found that whereas most control mitotic cells harbored two MTOCs, TRIM37 depletion resulted in an increased frequency of cells with more than two MTOCs, which often differed in size (Fig. 2A, Fig. S2A, Fig.2B). In addition, we found that ~40% of Cenpas did not nucleate microtubules, indicative of some composition heterogeneity (Fig. 2A, siTRIM37, inset 1). We conclude that microtubules nucleated from Cenpas contribute to the defective spindle assembly and chromosome miss-segregation phenotype of TRIM37 depleted cells.

**Figure 2.**
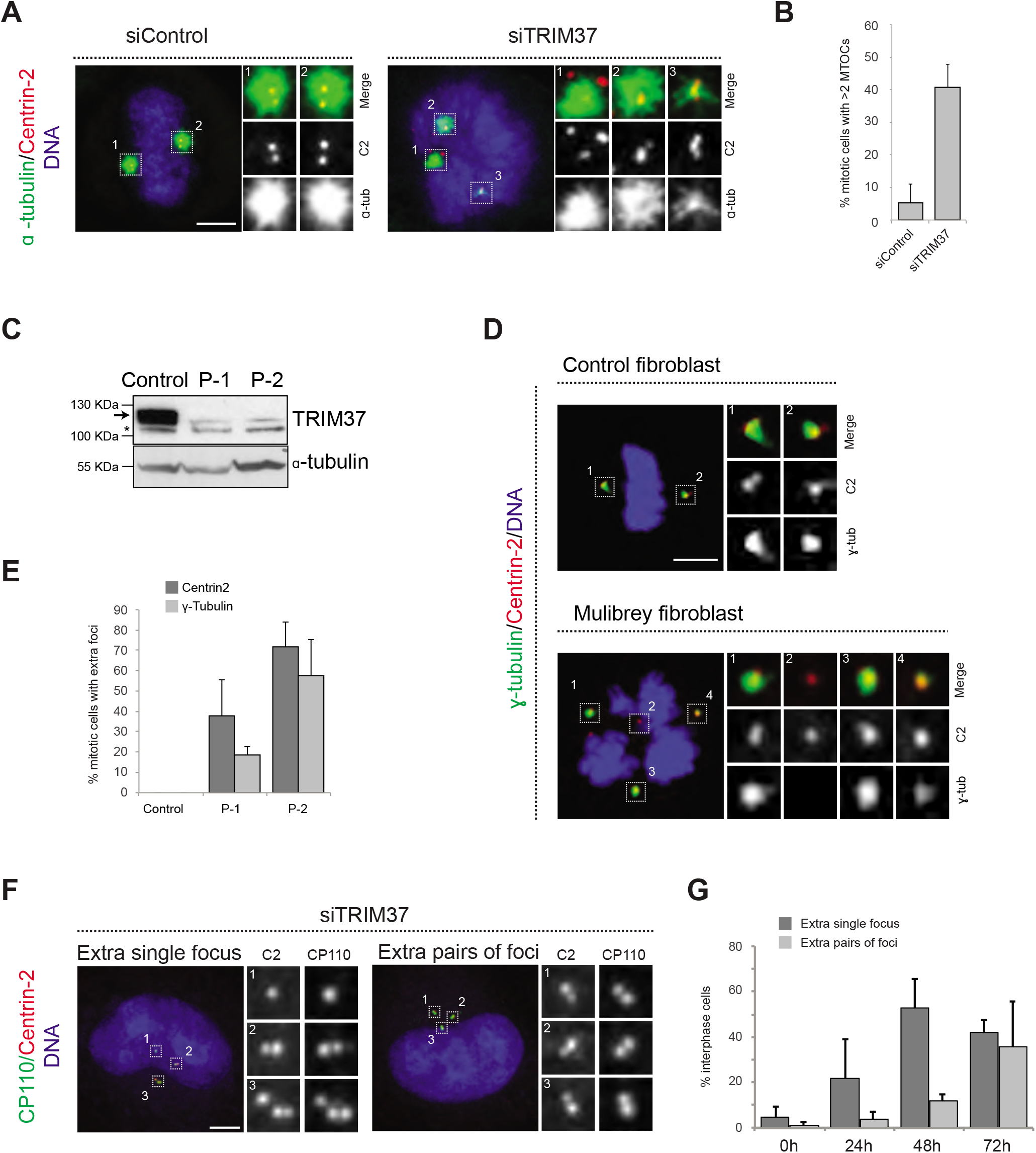
Cenpas can behave as extra MTOCs, including in Mulibery patient cells, and trigger new rounds of centriole duplication. **A.** Microtubule depolymerization-regrowth experiment in mitotic HeLa cells treated with control or TRIM37 siRNAs. Microtubules were depolymerized by a 30 min cold shock followed by 1-2 min at room temperature before fixation of cells and immunostaining for Centrin-2 and α-tubulin. **B.** Corresponding percentage of mitotic cells with >2 MTOCs; P<0.01. Note that ~40% of the extra Centrin-2 foci observed in mitosis did not nucleate microtubules, as is the case for two of them in inset 1 (siTRIM37). N= 40 Cenpas each scored in three independent experiments, SD 12.6%. **C.** Western blot of cell lysates from control and patient (P-1, P-2) fibroblasts probed with antibodies against TRIM37 (top) or α-tubulin as loading control (bottom). The arrow indicates TRIM37, the asterisk a non-specific band. Select molecular weight markers are indicated in kDa in this and other Western blot panels. **D.** Control and patient-1 (P-1) fibroblasts in mitosis immunostained for Centrin-2 and γ-tubulin. **E.** Corresponding percentage of mitotic cells with extra number of Centrin-2 or γ-tubulin foci in control and patient (P-1 and P-2) fibroblasts. Data from a total of 57 (P-1) and 54 (P-2) mitotic cells obtained from three independent experiments. **F.** HeLa cells depleted of TRIM37 and immunostained for Centrin-2 plus CP110, illustrating a case with an extra single focus (left, inset 1) and one with an extra pair of foci (right). **G.** Corresponding percentage of interphase cells with extra single focus or extra pairs of foci.

To further explore the importance of Cenpas, we addressed whether they are also present in Mulibrey patient cells. Using healthy donor fibroblasts as controls, we analyzed fibroblasts derived from two patients bearing the Finnish founder mutation, the most frequent TRIM37 disease alteration, which results in a frame shift of the coding sequence generating a premature stop codon (Avela et al., 2000). As shown in Figure 2C, Western blot analysis showed essentially no detectable TRIM37 protein in patient cells. We immunostained control and patients fibroblast with antibodies against Centrin-2 to monitor the presence of Cenpas, as well as against γ-tubulin to probe their ability to recruit PCM and, thereby, to nucleate microtubules. Echoing the results in tissue culture cells depleted of TRIM37, we found that patient cells in mitosis harbored supernumerary Centrin-2 foci, some of which were positive for γ-tubulin (Fig. 2D, 2E). Patient cells also exhibited evidence of chromosome miss-segregation, as would be expected from multipolar spindle assembly (Fig. 2D). We conclude that Cenpas are present and active also in Mulibrey patient cells.

We set out to address whether Cenpas in tissue culture cells are also active in triggering further rounds of centriole assembly, potentially in a subsequent cell cycle to the one in which they formed. To this end, we transfected cells with TRIM37 siRNAs and monitored the presence of Cenpas 24, 48 and 72 hours thereafter using antibodies against Centrin-2 and CP110. Control cells harbored two individual Centrin-2/CP110 foci in G1 and two pairs of such foci in S/G2, corresponding to two pairs of resident centriole plus procentriole (Fig. S2B). Upon TRIM37 depletion, we found that supernumerary Centrin-2/CP110 foci appeared principally as individual units at the 24 hours time point, but that pairs of foci became more frequent at the 48 and 72 hours time points (Fig. 2F, 2G). We conclude that Cenpas can trigger further rounds of centriole assembly.

### Ultra expansion microscopy and electron microscopy reveal aberrant centriole-related structures upon TRIM37 depletion

We set out to address whether Cenpas exhibit further hallmarks of centrioles. We thus tested whether Cenpas harbor microtubules characteristic of centrioles by staining cells depleted of TRIM37 with antibodies against acetylated tubulin, a signature modification of centriolar microtubules, finding that ~23% such cells possessed extra acetylated tubulin foci (Fig. 3A, 3B). To examine this feature at higher resolution, we analyzed cells using ultrastructure expansion microscopy (U-ExM) coupled to confocal imaging (Gambarotto et al., 2019). RPE-1 cells expressing Centrin1:GFP were immunostained for GFP to identify Cenpas, for CEP152 to mark mature centrioles and for acetylated tubulin. Control cells contained two mature centrioles positive for all three markers (Fig. 3C). We found that some Cenpas formed upon TRIM37 depletion harbored merely Centrin1:GFP, but neither acetylated tubulin or CEP152 (Fig. 3D-3F, yellow arrows). By contrast, other Cenpas were positive for all three markers (Fig. 3E-3G), with the acetylated tubulin signal being smaller than normal in some cases (Fig. 3E, 3F, white arrows). Moreover, some Cenpas appeared to have matured into entities with regular looking acetylated tubulin and CEP152 signals (Fig. 3G). Together, these findings support the notion that Cenpas are heterogeneous in nature with partially overlapping composition.

**Figure 3.**
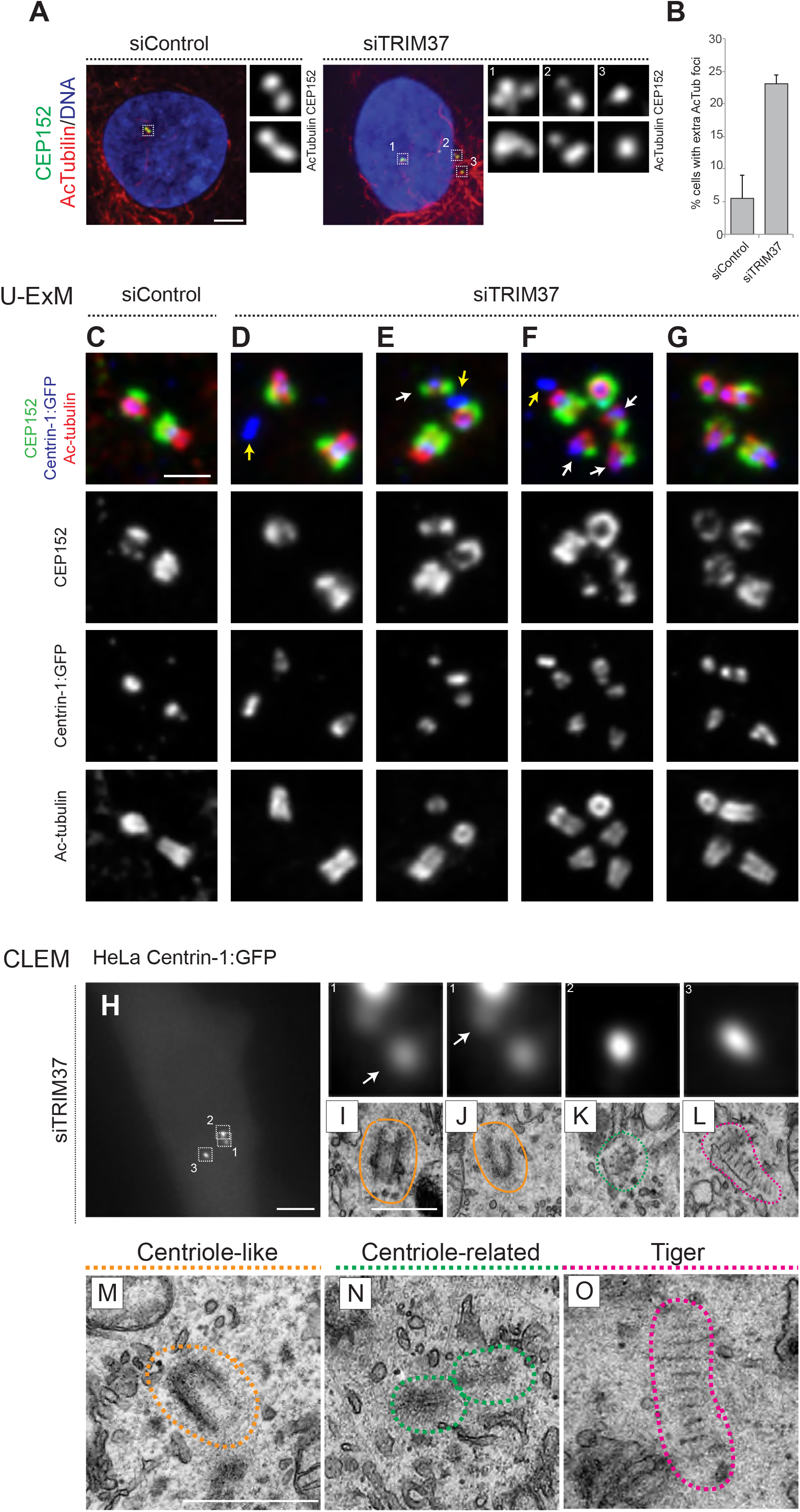
Cenpas are structures related to centrioles or electron-dense striped structures. **A.** RPE-1 cells treated with control or TRIM37 siRNAs, and immunostained for CEP152 plus acetylated tubulin. **B.** Corresponding percentage of cells with extra foci of acetylated tubulin; P<0.05. **C-G.** Ultrastructure expansion microscopy (U-ExM) confocal images of control (C) or TRIM37 (D-G) depleted RPE-1 cells expressing Centrin-1:GFP, and immunostained for GFP, CEP152 as well as acetylated tubulin. Yellow arrows point to Cenpas lacking CEP152 and acetylated tubulin, white arrows to those harboring both proteins, but with an unusual distribution. Scale bar 500 nm. **H-L.** CLEM analysis of HeLa cell (cell 3 in Fig. S3F) expressing Centrin-1:GFP and depleted of TRIM37. Maximal intensity projection of wide-field microcopy image covering the entire cell volume (H), and magnified insets from the light microscopy images above the corresponding EM images (I-L), with white arrows pointing to relevant Centrin-1:GFP focus. Scale bars: 5 μm in F, 500 nm in G. Here and in panels M-O, orange, green and pink dashed lines surround respectively centrioles-like, centriole-related and tiger structures. Filled orange lines surround resident centrioles. **M-O.** Centriole-like (M, cell 7 in Fig. S3F), centriole-related (N, cell 7 in Fig, S3F), and tiger (O, cell 2 in Fig, S3F) structures. Scale bar is 500 nm.

To uncover the ultrastructure of Cenpas, we conducted correlative light and electron microscopy (CLEM). Using fluorescence microscopy, we screened HeLa and RPE-1 cells expressing Centrin-1:GFP depleted of TRIM37 to identify Cenpas, using a gridded coverslip to acquire information regarding GFP foci position, before proceeding with serial section transmission electron microscopy (TEM). In addition to control cells (Fig. S3A-S3B), we analyzed 8 cells depleted of TRIM37 (Fig. 3H, Fig. S3C-F). From a total of 47 Centrin-1:GFP foci observed by light microscopy in TRIM37 depleted cells, serial section TEM analysis established that in addition to those corresponding to normal looking resident centrioles or procentrioles, 20 corresponded to unusual structures described hereafter (Fig. S3F). We found the expected number of resident centrioles (15 found/16 expected, see Fig. S3F; Fig. 3I, 3J), as well as two extra centriole-like structures in one cell (Fig. 3M; Fig. S3D). Eight of the other unusual structures were centriole-related electron-dense assemblies that harbored microtubules but only partially resembled centrioles (Fig. 3K, 3N; Fig. S3C, S3D). Strikingly, the remaining 12 other unusual structures were elongated electron-dense striped entities, hereafter referred as “tiger” structures (Fig. 3L, 3O; Fig. S3C, S3E). We noted also that an individual tiger structure sometimes correlated with more than one Centrin-1:GFP focus (Fig. S3E). Overall, we conclude that Cenpas forming upon TRIM37 depletion are heterogeneous in nature, only sometimes bearing resemblance to centrioles, perhaps reflecting different pathways or steps in their assembly.

### TRIM37 depletion triggers formation of elongated Centrobin assemblies

Because TRIM37 is an E3 ligase, the activity of which is important for preventing Cenpas formation (Balestra et al., 2013), we reasoned that a protein implicated in centriole assembly might accumulate in an aberrant manner upon TRIM37 depletion, causing the observed phenotype. Therefore, we conducted a small screen by immunostaining cells depleted of TRIM37 with antibodies against >20 centriolar and centrosomal proteins (Fig. S4A and data not shown). This analysis revealed that Centrobin, which normally localizes tightly to the daughter centriole and to procentrioles (Zou et al., 2005), is present in striking elongated cytoplasmic assemblies upon TRIM37 depletion (Fig. 4A, 4B). We found that ~80% of TRIM37 depleted cells bear usually one or two such Centrobin assemblies (Fig. S4B, S4C). Furthermore, SPICE, which is involved in centriole biogenesis, was also present in Centrobin assemblies upon TRIM37 depletion, even though SPICE was not needed for their formation (Fig. S4D, S4E). Remarkably, all cells with Cenpas were positive for Centrobin assemblies (n=150) with Cenpas often colocalizing with them (Fig. 4A, 4B). In summary, aberrant Centrobin assemblies are invariably present in cells with Cenpas and are often associated with them.

**Figure 4.**
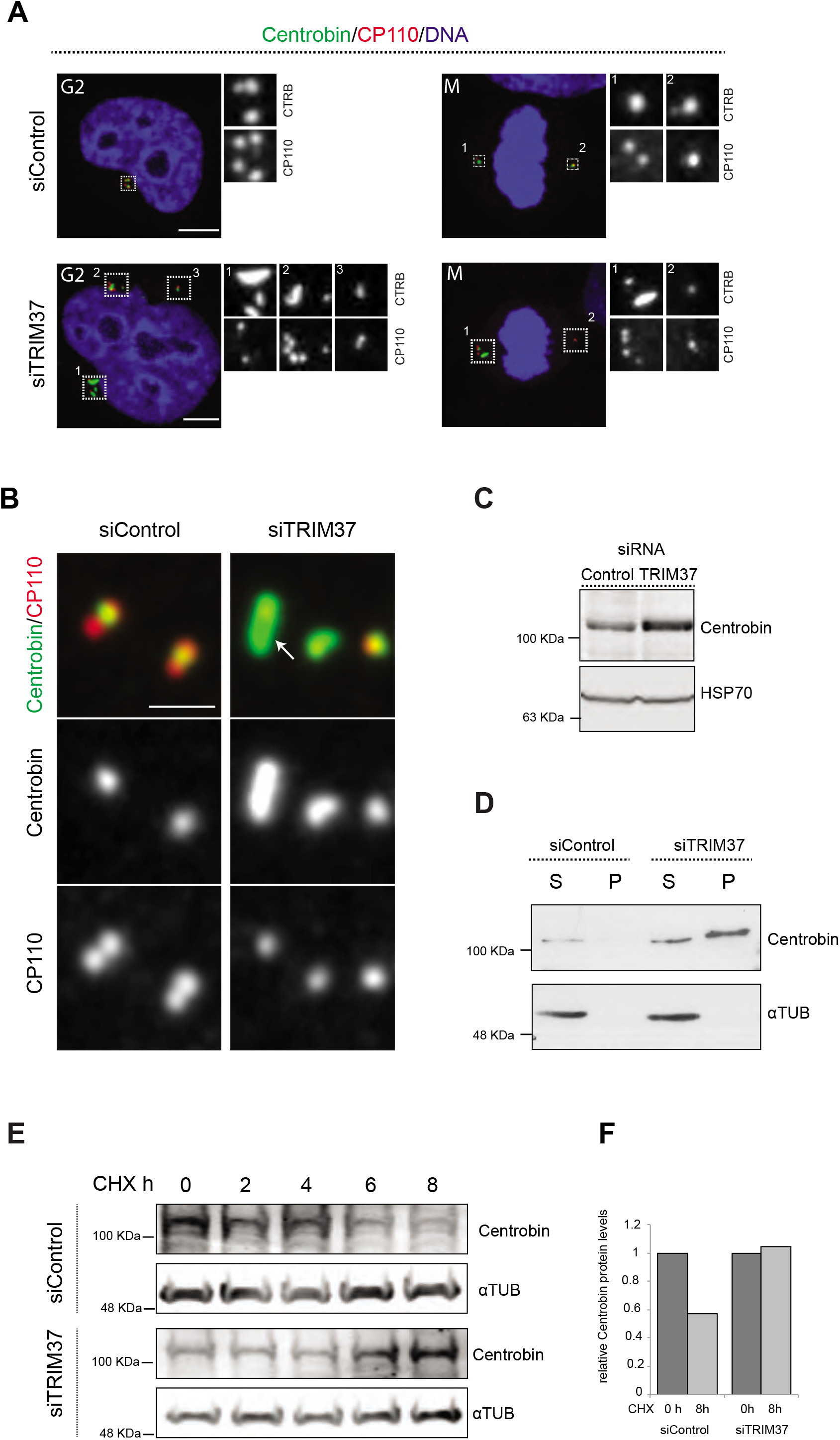
TRIM37 regulate Centrobin protein stability and levels. **A.** HeLa cells in G2 or mitosis, as indicated, treated with control or TRIM37 siRNAs, and immunostained for CP110 plus Centrobin. **B.** High magnification confocal view of cells treated with control or TRIM37 siRNAs immunostained for Centrobin and CP110. Arrow points to elongated Centrobin assembly. Scale bar 1 μm. **C.** Western blot of lysates from HeLa cells treated with control or TRIM37 siRNAs probed with antibodies against Centrobin (top) or HSP70 as loading control (bottom). **D.** Western blot of soluble (S) or insoluble (P, for pellet) fractions of lysates from HeLa cells treated with control or TRIM37 siRNAs, probed with antibodies against Centrobin (top) or α-tubulin as loading control (bottom). Note that Centrobin present in the insoluble fraction migrates slower upon TRIM37 depletion, suggestive of some posttranslational modification. **E.** Western blot of total Centrobin protein levels in control and TRIM37 depleted HeLa cells treated with cycloheximide (CHX) for indicated time in hours (h), probed with antibodies against Centrobin (top) or α-tubulin as loading control (bottom). Note that the amount of lysate loaded for the TRIM37 depleted sample was ~50% of that loaded for the siControl condition. **F.** Quantification of relative Centrobin protein levels from the Western blot shown in E.

How could TRIM37 regulate Centrobin? Performing real time quantitative PCR experiments showed a mere slight diminution in Centrobin mRNA levels upon TRIM37 depletion (Fig. S4F), suggesting that regulation is not at the transcriptional level. By contrast, Western blot analysis uncovered that Centrobin protein levels were increased upon TRIM37 depletion (Fig. 4C). Given the elongated Centrobin assemblies identified by immunostaining, we speculated that the overall increase in Centrobin protein level might reflect an accumulation into such structures, potentially in an insoluble form. Accordingly, fractionating cell lysates into soluble and insoluble fractions, we found that the increase in Centrobin protein levels was most pronounced in the latter (Fig. 4D). We noted also that the insoluble pool of Centrobin migrated slower in the gel upon TRIM37 depletion, suggesting that TRIM37 not only restricts Centrobin levels, but also regulates its posttranslational state in some manner.

Since TRIM37 is an E3 ubiquitin ligase, we reasoned that its activity might modulate Centrobin protein degradation and, thereby, stability. Therefore, we assayed the stability of the Centrobin protein pool over time in the presence of the translation inhibitor Cycloheximide. As reported in Figure 4E and 4F, we found that TRIM37 depletion significantly increased Centrobin protein stability. One possibility would be that TRIM37 ubiquitinates Centrobin, thus targeting it for degradation, such that increased Centrobin levels upon TRIM37 depletion would trigger formation of Centrobin assemblies and Cenpas. However, although Centrobin overexpression generates aggregates (Jeong et al., 2007), we found that such aggregates did not resemble the elongated Centrobin assemblies nor did they trigger Cenpas formation (Fig. S4G). In addition, TRIM37 overexpression did not alter Centrobin centrosomal distribution (Fig. S4H). Moreover, no evidence for TRIM37 mediated Centrobin ubiquitination was found in cell free assays (data not show), such that the detailed mechanisms of Centrobin modulation by TRIM37 remain to be deciphered. Regardless, we conclude that TRIM37 normally regulates Centrobin stability, preventing the protein from forming elongated assemblies invariably present in cells with Cenpas.

### Centrobin assemblies may serve as platforms for Cenpas formation

We set out to further characterize the elongated Centrobin assemblies formed upon TRIM37 depletion and assay their role in Cenpas generation. We used U-ExM coupled to STED super-resolution microscopy to analyze the distribution of Centrobin upon TRIM37 depletion at higher resolution. We immunostained RPE-1 cells expressing Centrin-1:GFP with antibodies against GFP, CEP152 and Centrobin. In control conditions, centrioles viewed in cross section exhibited a clear localization of Centrobin between the outer CEP152 and the inner Centrin-1:GFP signals (Fig. 5A). Cells depleted of TRIM37 exhibited analogous distributions at resident centrioles (Fig. 5A), but also harbored elongated Centrobin assemblies often abutting Centrin-1:GFP foci (Fig. 5A, arrows). Strikingly, the superior resolution afforded by U-ExM coupled to STED revealed that such Centrobin assemblies are striated (Fig. 5A). Suggestively, the inter-stripe distances of these Centrobin assemblies were analogous to those of the tiger structures unveiled through CLEM (Fig. 5B). In summary, U-ExM analysis strongly suggests that Centrobin is a constituent of the electron-dense tiger structures observed by TEM upon TRIM37 depletion, and raises the possibility that such structures serve as platforms for Cenpas formation.

**Figure 5.**
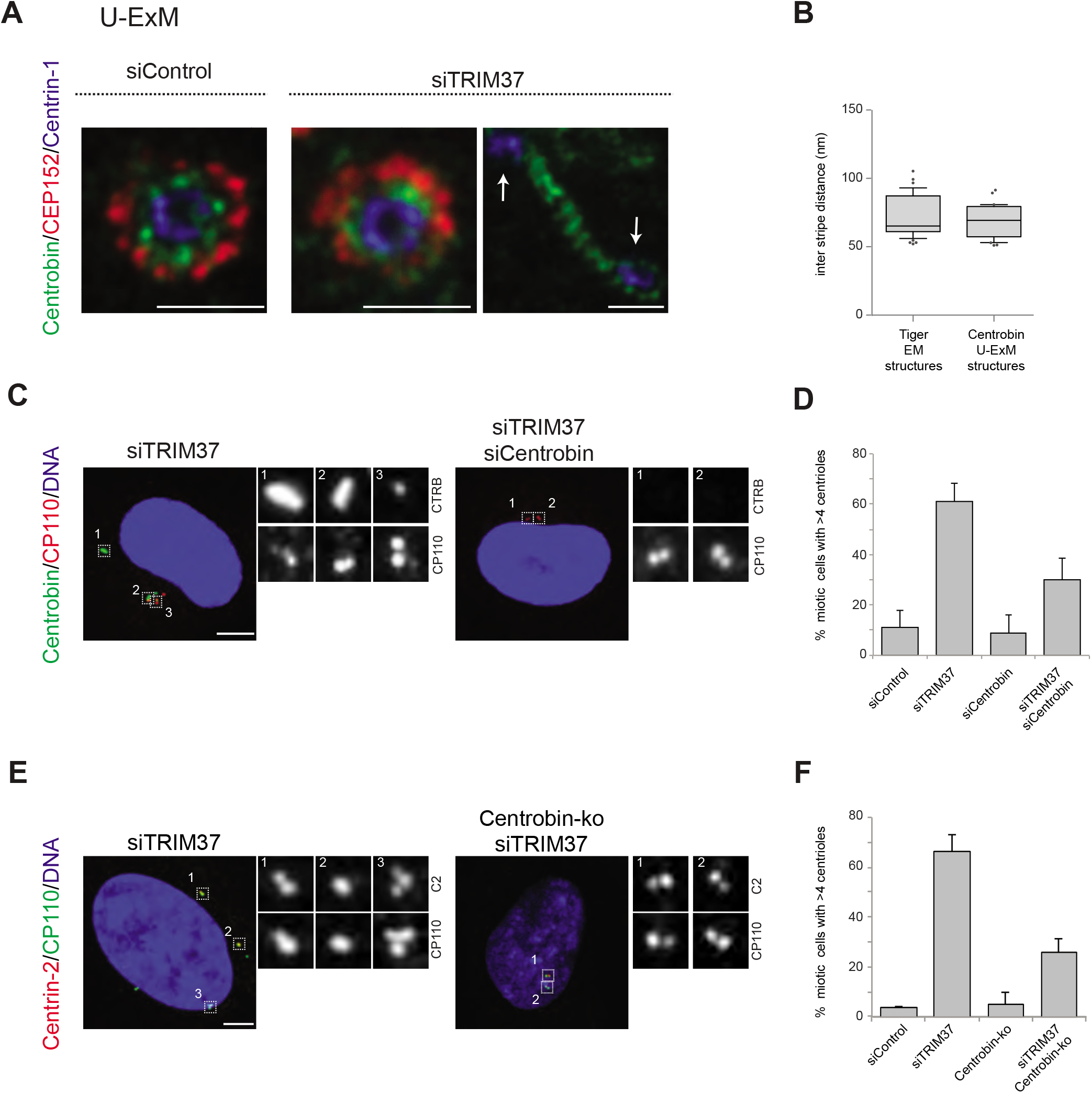
Centrobin promotes Cenpas formation. **A.** U-ExM coupled to STED super-resolution microscopy of RPE-1 cells immmunostained for CEP152, Centrin-1 and Centrobin. White arrows point to Cenpas in close proximity to Centrobin assembly. Scale bars are 250nm. **B.** Box-and-whisker plot of inter stripe distances in TEM tiger structures (n = 53 from 5 tiger structures) and U-ExM Centrobin structures (n = 30 from 3 Centrobin structures); P=0.38, not significant. **C.** HeLa cells depleted of TRIM37 alone or simultaneously of TRIM37 and Centrobin, and immunostained for CP110 plus Centrobin. **D.** Corresponding percentages of mitotic cells treated with the indicated siRNAs harboring >4 CP110 foci. siTRIM37 versus siTRIM37+siCentrobin: P < 0.05. **E.** Control and Centrobin-ko RPE-1 cells transfected with TRIM37 siRNAs immunostained for Centrin-2 and CP110. **F.** Corresponding percentages of mitotic cells with >4 CP110 foci. siTRIM37 versus siTRIM37+Centrobin-KO: P < 0.01.

To investigate the potential role of Centrobin in Cenpas formation, we tested whether Centrobin depletion reduces Cenpas numbers in cells depleted of TRIM37. Although Centrobin depletion was reported initially to impair centriole assembly in HeLa cells (Zou et al., 2005), more recent work with Centrobin knock out cells (Centrobin-ko) demonstrates that the protein is dispensable for this process in RPE-1 cells (Ogungbenro et al., 2018). In our hands, siRNA-mediated depletion of Centrobin did not impact centriole assembly either in HeLa Kyoto cells, despite near-complete protein depletion (Fig. S5A-C). As anticipated, Centrobin assemblies disappeared entirely from cells doubly depleted of Centrobin and TRIM37 (Fig. 5C). Importantly, we found that Cenpas number was significantly lowered in such doubly depleted cells compared to cells depleted of TRIM37 alone (Fig. 5D). Interestingly, however, even if Centrobin depletion was complete as judged by Western blot analysis (Fig. S5C), Cenpas formation upon TRIM37 depletion was only partially prevented by Centrobin siRNA treatment (Fig. 5D). To test whether this might have been due to residual Centrobin or TRIM37 in the double siRNA depletion setting, we performed a similar experiment with RPE-1 Centrobin-ko cells (Ogungbenro et al., 2018), reaching similar conclusions (Fig. 5E, 5F, Fig. S5D). Together, these results support the view that upon TRIM37 depletion Centrobin assemblies act as platform seeding the formation of some, but not all, Cenpas.

### Centrobin and PLK1 together promote Cenpas assembly upon TRIM37 depletion

To further understand the mechanisms of Cenpas formation upon TRIM37 depletion, we tested if select centriolar proteins that are critical for canonical centriole duplication were also needed for Cenpas generation. To test the role of PLK4, HeLa cells were grown in the presence of Centrinone for 5 days and then depleted of TRIM37 for 3 days in the continued presence of Centrinone. We found that Cenpas did not form under these conditions, demonstrating an essential role for PLK4 kinase activity (Fig. 6A, 6B). We also tested the requirement for HsSAS-6, STIL, CPAP and SPICE. As anticipated, single depletion of these components resulted in decreased centriole number (Fig. 6A). However, depletion of STIL, CPAP or SPICE did not dramatically modify the number of Cenpas upon TRIM37 depletion (Fig. 6A). By contrast, HsSAS-6 depletion greatly reduced Cenpas number (Fig. 6A). To further explore the impact of HsSAS-6, we depleted TRIM37 from RPE-1 p53-/- HsSAS-6 knock out cells (HsSAS-6-ko) (Wang et al., 2015). Although HsSAS-6-ko cells invariably lacked centrioles (Fig. 6A, 6B), some Cenpas nevertheless formed upon TRIM37 depletion, although to a lesser extent than following depletion of TRIM37 alone (Fig. 6A, 6B). Moreover, we found that elongated Centrobin assemblies were generated unabated upon TRIM37 depletion in cells treated with Centrinone or lacking HsSAS-6 (Fig. 6C; Fig. S6). We conclude that PLK4 and HsSAS-6 act downstream of Centrobin in the pathways leading to Cenpas formation upon TRIM37 depletion.

**Figure 6.**
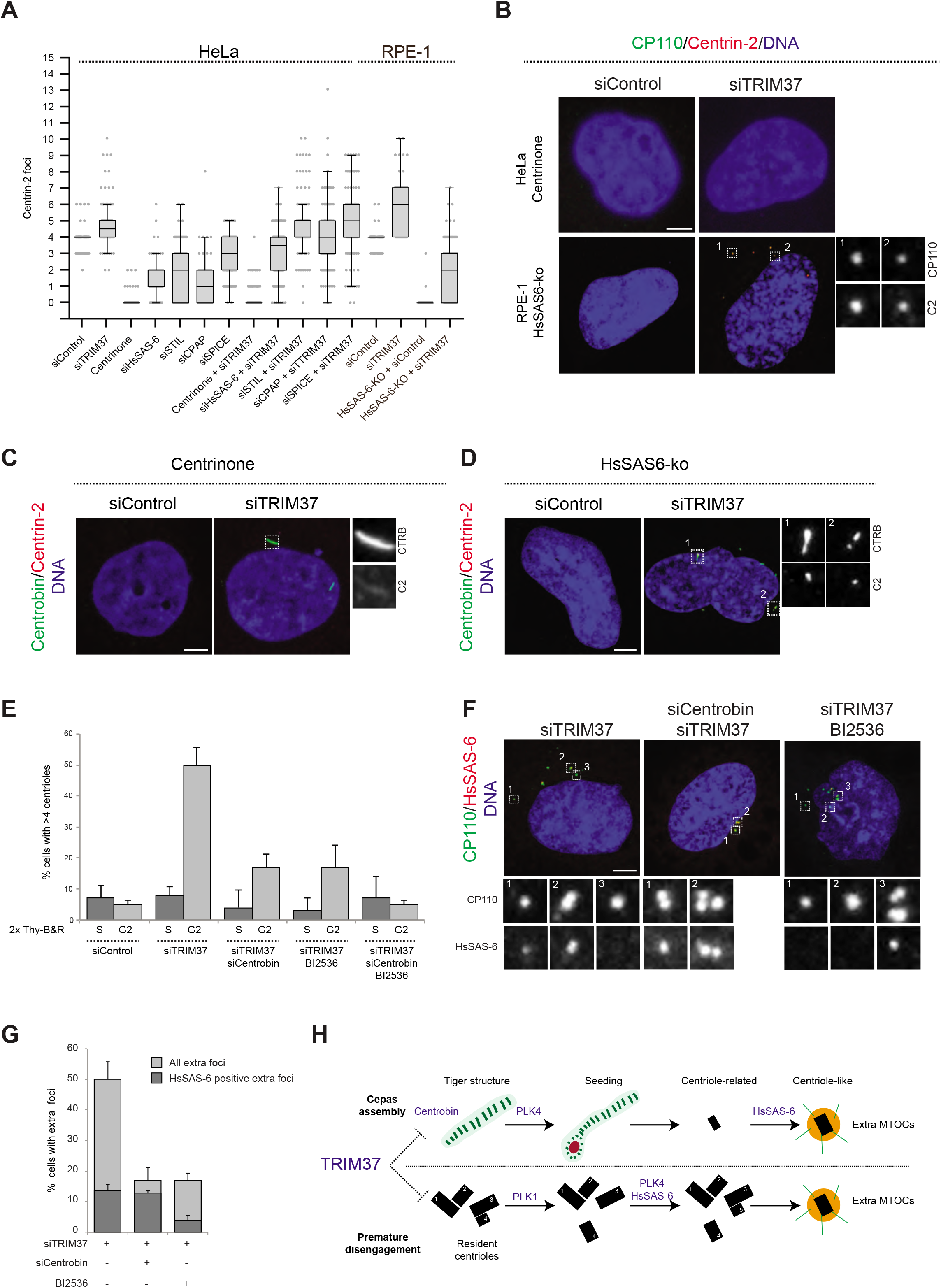
Two pathways contribute to Cenpas formation upon TRIM37 depletion. **A.** Box-and-whisker Tukey plot of Centrin-2 foci number per cell in indicated conditions. All cells were analyzed in mitosis with the exception of HsSAS-6-ko conditions. Centrinone versus centrinone + siTRIM37: P < 0.01, HsSAS-6-ko vs HsSAS-6 ko + siTRIM37: P < 0.0001. **B.** HeLa cells grown with centrinone for 8 days (top) or RPE-1 HsSAS-6-ko cells (bottom), both treated with control or TRIM37 siRNA, before immunostaining for CP110 and Centrin-2 and CP110. **C-D.** HeLa cells grown with centrinone for 8 days (C) or RPE-1 HsSAS-6-ko cells (D), both treated with control or TRIM37 siRNA, before immunostaining for Centrobin and Centrin-2. **E.** HeLa cells were synchronized with a double thymidine block, released and transfected with control, TRIM37, Centrobin, or both TRIM37 and Centrobin siRNAs, as indicated. Additionally, DMSO or BI-2536 was added to the cells, which were fixed at time 0 h or 8vh after release, before immunostaining with antibodies against CP110 and Centrobin. The percentage of cells with extra CP110 foci was quantified in each condition. siTRIM37+siCentrobin versus siTRIM37+siCentrobin+BI2536: P = 0.063, non significant. **F.** HeLa cells were synchronized with a double thymidine block, released and transfected with control, TRIM37, Centrobin, or both TRIM37 and Centrobin siRNAs, as indicated. Additionally, DMSO or BI-2536 was added to the cells, which were fixed at time 0 h or 8 h after release, before immunostaining with antibodies against CP110 and HsSAS-6. **G.** Corresponding percentage of cells with extra CP110 foci, with an indication of the fraction of them bearing HsSAS-6. The percentage of cells bearing extra HsSAS-6 foci between siTRIM37+siCentrobin versus siTRIM37+BI2536 was significant: P value < 0.05. **H.** Working model of TRIM37 role in preventing formation of supernumerary MTOCs. Our findings lead us to propose that TRIM37 prevents the formation of supernumerary Centrin foci through two independent pathways mediated by Centrobin (top) and PLK 1 (bottom). The Centrobin pathway relies on the assembly of tiger Centrobin assemblies that act as platforms for PLK4-dependent Cenpas assembly. Thereafter, Cenpas could evolve into centriole-related and then centriole-like structures with the stepwise incorporation of other centriolar proteins such as HsSAS-6. We propose that the PLK1 pathway might reflect its role in promoting centriole disengagement. Note that only extra MTOCs are represented. See text for details.

To further uncover requirements for Cenpas generation, considering that PLK1 contributes only partially to their formation (Balestra et al., 2013), and that we found here the same to be true for Centrobin, we set out to investigate whether the combined removal of PLK1 and Centrobin may fully prevent Cenpas generation. To avoid the negative impact of PLK1 inhibition on cell cycle progression, we performed these experiments in synchronized cells depleted of TRIM37, and monitor Cenpas appearance during G2 after release from an S phase arrest. These cells were also subjected to Centrobin depletion and/or BI-2536 treatment to inhibit PLK1. Importantly, we found that simultaneous Centrobin depletion and PLK1 inhibition completely prevented Cenpas formation (Fig. 6E), indicating that PLK1 and Centrobin act in parallel to promote Cenpas formation upon TRIM37 depletion.

Further evidence supporting the existence of two parallel pathways towards Cenpas generation was obtained by examining the distribution of HsSAS-6 in cells depleted of TRIM37 plus either PLK1 or Centrobin. Indeed, we found that Cenpas generated upon combined TRIM37 depletion and PLK1 inhibition, which thus rely strictly on Centrobin, rarely harbored HsSAS-6 (Fig. 6F, 6G). By contrast, Cenpas generated upon double depletion of TRIM37 and Centrobin, which thus rely strictly on PKL1, frequently harbored HsSAS-6 (Fig. 6F, 6G). Taken together, our findings indicate that two pathways are triggered when TRIM37 is lacking: one that relies on Centrobin assemblies that act as a platform to assemble Cenpas, which at the least is initially independent of HsSAS-6, and another one mediated by PLK1 that operates through HsSAS-6 recruitment (Fig. 6G, see discussion).

## DISCUSSION

Centriole number control is critical for proper cell physiology, including genome integrity. Assemblies of centriolar proteins that can recruit PCM and nucleate microtubules despite not being *bona fide* centrioles must likewise be kept in check. Here, we identify the TRIM37 E3 ligase, which is mutated in Mulibrey nanism, as a critical component that prevents the formation of centriolar protein assemblies (Cenpas) through two parallel pathways relying on PLK1 and Centrobin. Of particular interest, we uncover that TRIM37 regulates the stability of Centrobin, which upon TRIM37 depletion forms striated structures that we propose serve as platforms for Cenpas generation.

### Two pathways together result in Cenpas upon TRIM37 depletion

What are the mechanisms leading to Cenpas formation upon TRIM37 depletion? We previously hypothesized that TRIM37 could act by restricting centriole reduplication in G2, since PLK1 inhibition in TRIM37 depleted cells reduced Cenpas formation (Balestra et al., 2013). However, although blocking PLK1 activity in TRIM37 depleted cells reduced Cenpas numbers, some remained despite such inhibition (Balestra et al., 2013). Moreover, while Cenpas form upon TRIM37 depletion as early as 4 h after the G1/S transition (Balestra et al., 2013), PLK1-mediated centriole reduplication occurs only 24 h after G2 arrest (Loncarek et al., 2010). It may even be that the role exerted by PLK1 following TRIM37 depletion is not linked to its known function in regulating licensing. Regardless, we obtained further evidence here that Cenpas do not form solely through a reduplication mechanism. First, some Cenpas appear away from resident centrioles. Second, most Cenpas do not harbor the procentriolar protein HsSAS-6, at the least initially, which is in contrast to the situation during centriole reduplication during G2 arrest. Furthermore, analysis with CLEM revealed that Cenpas are usually either centriole-related structures or novel striped electron-dense structures. Together, these findings indicate that Cenpas do not form solely through centriole reduplication, but also through an alternative novel *de novo* pathway.

Our findings indicate that this alternative pathway relies on Centrobin: whereas the sole removal of Centrobin also merely decreases Cenpas number, the joint removal of PLK1 and Centrobin entirely prevent their generation (Fig. 6E).

The heterogeneity in Cenpas ultrastructure uncovered by CLEM might also reflect the co-existence of these independent assembly pathways. Such heterogeneity may reflect in addition a step-wise nature of the *de novo* generation process. This possibility is in line with the fact that more extra HsSAS-6 foci are present 72 hours after transfection with TRIM37 siRNAs (Balestra et al., 2013), compared to the 48 hours post-transfection analyzed here (Fig. 1C). Therefore, HsSAS-6 might not be present or required for the onset of *de novo* Cenpas formatoin, but could contribute later to their consolidation. In line with this view, HsSAS-6-k-o cells depleted of TRIM37 can assemble some Cenpas, perhaps more rudimentary ones. Interestingly in addition, this observation further suggests that *de novo* Cenpas generation upon TRIM37 depletion must in some way differ from the classical *de novo* centriole assembly, which is fully reliant on HsSAS-6 (Wang et al., 2015).

We found that one protein that is essential for forming all Cenpas upon TRIM37 depletion is PLK4, which is also required for centriole reduplication and *de novo* centriole assembly following Centrinone treatment (Habedanck et al., 2005; Wong et al., 2015). How could PLK4 be required for Cenpas generation stemming from the Centrobin assemblies formed upon TRIM37 depletion? PLK4 condensates forming away from resident centrioles have been observed in RPE-1 TRIM37 knock out (TRIM37-ko) cells (Meitinger et al., 2016). However, we did not detect such PLK4 localization, perhaps reflecting differences between chronic versus acute TRIM37 depletion. Another difference potentially related to distinct depletion regimes is that upon Centrinone treatment, TRIM37-ko cells form centrosome-like structures harboring notably PLK4 and HsSAS-6, and which recruit PCM components, behaving as MTOCs (Meitinger et al., 2016). This is in contrast to our findings whereby no Cenpas forms upon treatment with TRIM37 siRNAs and Centrinone. Regardless, it is interesting to note that in *Xenopus* extracts, PLK4 self-assembles into condensates that recruit γ-tubulin and behave as MTOCs (Montenegro Gouveia et al., 2018), raising the possibility that Centrobin assemblies may serve as platforms to recruit such condensates.

### Centrobin as a TRIM37 target

What are the targets of TRIM37 that are relevant for restricting Cenpas formation? Our work suggests that a critical target is Centrobin, since upon TRIM37 depletion Centrobin protein levels increase and elongated Centrobin assemblies form in the cytoplasm. U-ExM coupled to STED super-resolution microscopy reveals that these Centrobin assemblies have a striped pattern akin to the structures uncovered by CLEM, and are often intimately linked with Cenpas. Given the cytoplasmic localization of these assemblies, this role of TRIM37 is likely to be exerted by the cytoplasmic protein pool rather than the centrosomal or the nuclear ones. Centrobin contributes to several aspects of centriole assembly and growth, as well as ciliogenesis (Gudi et al., 2011; Ogungbenro et al., 2018; Zou et al., 2005). These functions might be linked to Centrobin’s ability to stabilize and promote microtubule nucleation (Gudi et al., 2011; Jeong et al., 2007; Shin et al., 2015). Future work will undoubtedly clarify the molecular connection between TRIM37 and Centrobin. Because Centrobin depletion does not fully prevent Cenpas formation upon TRIM37 depletion, other TRIM37 targets must be invoked, and PLK1 or a protein regulating its activity, is an attractive possibility in this respect.

### Cenpas form through different routes but similarly threaten cell physiology

Although with different molecular origins, centriolar protein assemblies have been reported in other contexts (Li et al., 2012; Shiratsuchi et al., 2015). Thus, the centriolar protein Neurl4 interacts with CP110 and promotes its destabilization, such that Neurl4 depletion results in increased CP110 protein levels and formation of ectopic MTOCs (Li et al., 2012). Likewise, depletion of RMB14 triggers the formation of centriolar protein complexes that do not initially require HsSAS-6 for their assembly. RBM14 normally limits formation of the STIL/CPAP complex, which upon RMB14 depletion triggers aberrant centriolar protein complex formation (Shiratsuchi et al., 2015). In both cases, however, Centrobin distribution was inspected and no elongated structures as the ones reported here were observed (Li et al., 2012; Shiratsuchi et al., 2015), suggesting different assembly routes. Although these previously reported centriolar protein assemblies and the ones analyzed here do not share a clear common molecular composition or assembly pathway, we propose to group them jointly under the acronym Cenpas, reflecting the fact that they similarly form following a *de novo* process and entail centriole-related structures that behave as active MTOCs.

To our knowledge, our work is the first example in which Cenpas have been reported in a human genetic disorder. The fact that Cenpas are present in Mulibrey derived patient cells raises the possibility that some disease features could be due to Cenpas formation, perhaps owing to the extra MTOCs and resulting chromosome miss-segregation phenotype. As one of the characteristics of Mulibrey nanism is the propensity to develop tumors, we speculate that the presence of Cenpas could contribute to this phenotype since extra centrioles can promote tumorigenesis (Ganem et al., 2009; Godinho et al., 2014; Levine et al., 2017; Sercin et al., 2016). We further speculate that some of the instances in which extra centriole numbers are observed in solid and hematological tumors may in reality correspond to Cenpas.

## ACKNOWLEDGMENTS

We thank Graham Knott and Marie Croisier (BioEM platform of the School of Life Sciences, EPFL) for assistance with CLEM and TEM, Isabelle Flückiger for technical support, as well as Niccolò Banterle for advice with U-ExM. Anna-Elina Lehesjoki (Department of Medical and Clinical Genetics, University of Helsinki, Finland) is acknowledged for her contribution towards securing patient and control fibroblast, Daniel Gerlich (Vienna BioCenter, Austria), Ciaran Morrison (Centre for Chromosome Biology, Galway, Ireland), Bryan Tsou (Memorial Sloan Kettering Cancer Center, New York, US) and George Hatzopoulos (EPFL, Lausanne, Switzerland) for cell lines, Michel Bornens (Institut Curie, Paris, France), Andrew Holland (Johns Hopkins University School of Medicine, Baltimore, USA) and Laurence Pelletier (Lunenfeld-Tanenbaum Research Institute, Toronto, Canada) for antibodies. We are also grateful to Niccolò Banterle and Fernando Monje Casas for critical reading of the manuscript. This work has been supported in part by the Swiss Cancer league (KFS-3388-02-2014, to P.G.). A.B. was supported by the National Centre for Competence in Research (NCCR) Chemical Biology, funded by the Swiss National Science Foundation. FRB thanks PG for having hosted and supported him at the beginning of the project. F.R.B. was funded by a Marie Curie IEF Postdoctoral Fellowship (PIEF_GA-2013-629414; CSIC-Cabimer, Seville, Spain) and by the University of Seville through a postdoctoral contract of the V PPIT-US (US-Cabimer, Seville, Spain). CABIMER is supported by the regional government of Andalucia (Junta de Andalucía).

## MATERIAL AND METHODS

### Cell culture, cell lines and cell treatments

HeLa Kyoto (generous gift from Daniel Gerlich) and U2OS (ATCC) cells were grown in high glucose DMEM medium (Sigma-Aldrich), hTERT-RPE-1(ATCC) cells in high glucose DMEM/F-12 medium (Sigma-Aldrich). Fibroblast cultures were established from skin biopsy samples of two Mulibrey nanism patients homozygous for the Finnish founder mutation, as well as a control individual, with approval by the Institutional Review Board of the Helsinki University Central Hospital (183/13/03/03/2009). The patients signed an informed consent for the use of fibroblast cultures. Fibroblasts were grown in RPMI medium (Sigma-Aldrich). Other cell lines used were HeLa cells carrying an integrated plasmid expressing Centrin-1:GFP (Piel et al., 2000), RPE-1 p53 -/- cells carrying an integrated plasmid (pCW57.1) expressing Centrin-1:eGFP under a doxycycline inducible promoter (generous gift from George Hatzopoulos), RPE-1 p53 -/- Centrobin knock out cells (Ogungbenro et al., 2018) (generous gift from Ciaran Morrison), and RPE-1 p53 -/- HsSAS-6 knock out cells (Wang et al., 2015) (generous gift from Bryan Tsou). All media were supplemented with 10% fetal bovine serum, 2 mM L-glutamine, 100 units/ml penicillin and 100 *μ* g/ml streptomycin (all from Sigma-Aldrich) and grown at 37°C in 5% CO2. HeLa Kyoto cells were synchronized using a double-thymidine block and release protocol as follows: cells were incubated in medium with 2 mM thymidine (Sigma Aldrich, T9250) for 17 h, released for 8 h and again incubated with 2 mM Thymidine for 17 h. For single transfection experiments, control or TRIM37 siRNAs transfections were performed during the 8 h period between the two thymidine treatments. For double transfection experiments, in addition to the above, either control or Centrobin siRNAs were transfected before the first Thymidine treatment. Drugs used in this work were 10 μM BI-2536 (S1109, Selleck Chemicals), 10 μM RO-3306 (Sigma-Aldrich, SML0569), 125 nM Centrinone (MCE, Hy-18682) and 150 μg/ml Cycloheximide (Sigma-Aldrich, C7698).

### Transfections, plasmids and siRNAs

For siRNA treatments, cells were typically transfected in a 6 well plate format with 20 μM siRNAs and 4 μL Lipofectamine RNAiMAX (Thermo Fisher Scientific); the depletion phenotype was inspected 72 h after transfection unless otherwise indicated in the text or the legends. siRNAs sequences were as follows: TRIM37 (5’-UUAAGGACCGGA GCAGUAUAGAAAA-3’) (Balestra et al., 2013), Centrobin (5’-AGUGCCAGACUGCAGCAACGGGAAA-3’) (Zou et al., 2005), SPICE (5’-GCAGCUGAGAACAAAUGAGUCAUUA-3’) (Archinti et al., 2010), HsSAS6 (5’-GCACGUUAAUCAGCUACAAUU-3’) (Strnad et al., 2007), STIL (5’-AACGUUUACCAUACAAAGAAA-3’) (Kitagawa et al., 2011a), CPAP (5’-AGAAUUAGCUCGAAUAGAA-3’) (Kitagawa et al., 2011a), and Stealth RNAi™ siRNA Negative Control Lo GC (Ref: 12935200; Invitrogen). For plasmid transient transfection, FuGENE 6 Transfection Reagent (Promega) was used according to the manufacturer’s protocol and the phenotype inspected 24 or 48 h after transfection. Transfected plasmids were as follows: pEBTet-TRIM37:GFP (Balestra et al., 2013), pGFP-Centrobin:GFP (pGFP-NIP2) (Shin et al., 2015) (generous gift from Kunsoo Rhee, Seoul National University, Korea) pEGFP:SPICE (Archinti et al., 2010) (generous gift from Jens Lüders, IRB, Barcelona, Spain) pcDNA3-TRIM37:GFP and pcDNA3-TRIM37:NES:GFP were generated by cloning the TRIM37 ORF (964 aa) fused to GFP or to the HIV-Rev NES sequence (LQLPPLERLTLD) (Wen et al., 1995) and GFP.

### Immunoblotting and Cycloheximide chase assay

For Western blot analysis, cells were lysed either in 2× Laemmli buffer (4% SDS, 20% glycerol, 125 mM Tris-HCl, pH 6.8) and passed 10 times through a 0.5 mm needle-mounted syringe to reduce viscosity, or in NP40 lysis buffer [10 mM Tris/HCl (pH 7.4)/150 mM NaCl/10% (v/v) glycerol/1% (v/v) Nonidet P40/1 mM PMSF and 1 μg/ml of each pepstatin, leupeptin, and aprotinin (Sigma-Aldrich) for 20 min at 4°C and then for 3 min at 37°C, before centrifugation at 20,000 g for 20 min. In this manner, the soluble fraction was separated from the insoluble pellet, which was then solubilized in 1x Laemmli buffer. Protein concentration was determined with a NanoDrop Spectrophotometer. Lysates were resolved by SDS-PAGE on a 10% polyacrylamide gel and immunoblotted on Immobilon-P transfer membrane (IPVH00010; 21 Millipore Corporation). Membranes were first blocked with TBS containing 0.05% Tween-20 (TBST) and 5% non-fat dry milk (TBST-5% milk) for 1h at room temperature, and then incubated with primary antibodies diluted in PBST-5% milk. Primary antibodies were 1:1000 rabbit anti-TRIM37 (A301-174A; Bethyl Laboratories), 1:30,000 mouse anti- *α* -tubulin (DM1a; Sigma-Aldrich), 1:500 rabbit anti-Centrobin (HPA023321; Atlas), and 1:20,000 anti-HSP70 (sc-24; Santa Cruz). Membranes were washed and incubated for 1h in secondary antibodies prepared also in TBST-5% milk. Secondary antibodies were 1:5,000 HRP-conjugated anti-rabbit (W4011; Promega) or mouse (W4021; Promega) IgGs. The signal was detected by standard chemiluminescence (34077; Thermo Scientific). Alternatively, polyacrylamide gels were immunoblotted on low fluorescence PVDF membranes (Immobilon-FL, Millipore), membranes blocked with Odyssey Blocking Buffer (LI-COR) and blotted with appropriate primary antibodies and 1:5,000 secondary antibodies IRDye 680RD anti-mouse IgG (H+L) Goat LI-COR (926-68070) and IRDye 800CW anti-rabbit IgG (H+L) Goat LI-COR (926-32211). Membranes were then air-dried in the dark and scanned in an Odyssey Infrared Imaging System (LI-COR), and images analyzed with ImageStudio software (LI-COR). In all cases, membrane washes were in TBST. For the cycloheximide chase experiment, HeLa Kyoto cells were treated with fresh DMEM containing 150 μg/ml cycloheximide (CHX). Cells were collected 0, 2, 4, 6 and 8 h after CHX addition, and protein extracts prepared in 2× Laemmli buffer as described above. 40 μg of siControl lysate and 20 μg of siTRIM37 lysate were resolved by SDS-PAGE, analyzed by immunoblotting with Centrobin and α-tubulin antibodies before quantification with ImageStudio. The siControl and siTRIM37 conditions at time 0 were normalized as 100%, and the other conditions for the same siRNA treatment expressed relative to this. Centrobin expression was quantified as the Centrobin signal divided by the α-tubulin signal.

### RNA isolation, reverse transcription and real-time PCR

RNA was extracted using the RNeasy Mini kit according to the manufacturer’s instruction (QIAGEN), including DNase I to avoid potential contaminations with DNA. 3 μg of total RNA, random hexamers and SuperScript III Reverse Transcriptase (InvitrogenTM) were used to obtain complementary DNA (cDNA). Quantitative PCR from cDNA was performed to assess siRNA-mediated knock-down of TRIM37 and Centrobin, using iTaq Universal SYBR Green Supermix following the manufacturer’s instructions (Bio-Rad) in an Applied Biosystems 7500 Fast Real-time PCR System (Thermo Fisher Scientific). Relative mRNA levels of the indicated genes were calculated by the 2-DDCT method (Bulletin 5279, Real-Time PCR Applications Guide, Bio-Rad), using GAPDH expression as endogenous control. The primer sequences used were: Centrobin:_CNTROB-FW 5’-GTCTCCATCTAGCTCAGCCC-3’, CNTROB-RV 5’-AGGCTCTGAATATGGCGCT C-3’, TRIM37: TRIM37-FW 5’-TGCCATCTTACGATTCAGCTAC-3’, TRIM37-RV 5’-CGCACAACTCCATTTCCATC-3’. GAPDH: GAPDH-FW 5’-GGAAGGTGAAGGTCGGAGTC-3’, GAPDH-RV 5’-GTTGAGGTCAATGAAGGGGTC-3’

### Indirect immunofluorescence and microtubule-regrowth assay

Cells were grown on glass coverslips and fixed for 7 min in −20°C methanol, washed in PBS, and blocked for 30 min in PBS 0.05% Tween 20 (PBST) with 1% bovine serum albumin. Cells were incubated overnight at 4°C with primary antibodies, washed three times for 5 min with PBST, incubated for 1 h at room temperature with secondary antibodies, washed three times for 5 min in PBST and mounted in Vectashield mounting medium with DAPI (H-1200; Vector Laboratories). Primary antibodies used for immunofluorescence were: 1:50 human anti-GFP from the recombinant antibody platform of Institut Curie (hVHH antiGFP-hFc, A-R-H#11), 1:1000 rabbit anti-GFP (RGFP-45ALY-Z; ICL), 1:500 mouse anti-HsSAS-6 (sc-81431; Santa Cruz), 1:1000 rabbit anti-CEP63 (06-1292; Millipore), 1:2000 rabbit anti-CEP152 (HPA039408; Sigma-Aldrich), 1:1000 mouse anti-acetylated tubulin (T6793; Sigma-Aldrich), 1:1000 mouse anti-γ-tubulin (GTU88, T5326; Sigma-Aldrich), 1:1000 mouse anti-Centrin2 (20H5; Sigma-Aldrich), 1:2000 rabbit anti-CEP164 (45330002; Novus Biologicals), 1;1000 mouse anti-α-tubulin (T6199; Sigma-Aldrich), 1:1000 rabbit anti-CP110 (12780-1-AP; Proteintech), 1:1000 mouse anti-Centrobin (ab70448; Abcam), 1:1000 rabbit anti-Centrobin (HPA023321; Atlas Antibodies), 1:1000 rabbit anti-CEP135 (ab75005; Abcam), 1:500 rabbit anti-CPAP (Kohlmaier et al., 2009), 1:500 rabbit anti-SPICE (HPA064843, Sigma-Aldrich), 1:8,000 rabbit anti-Ninein (L77), 1:1000 rabbit anti-hPOC5 (Azimzadeh et al., 2009) and 1:1000 rabbit anti-PLK4(KD) (Sillibourne et al., 2010) (both generous gifts from Michel Bornens, Institut Curie, Paris, France), 1:1000 rabbit anti-P-PLK4 (Moyer and Holland, 2019) (generous gifts from Andrew Holland), 1:400 mouse anti-C-Nap (611374; BD Biosciences) 1:2000 rabbit anti-STIL (ab222838; Abcam), 1:1000 rabbit anti-PCNT (ab4448; Abcam), 1:1000 mouse anti-AKAP450 (611518; BD Biosciences), 1:1000 rabbit anti-CDK5Rap2 (06-1398; Millipore), 1:1000 rabbit anti-CEP192 (a generous gift from Laurence Pelletier), 1:1000 rabbit anti-CEP170 (HPA042151; Sigma-Aldrich), 1:1000 mouse anti-P-T210-PLK1 (558400; BD Bioscience). Secondary antibodies were 1:500 mouse Alexa-488, 1:3000 rabbit Cy3, 1:3000 human Alexa-633, 1:1000 mouse Alexa-649, and 1:500 human Alexa-488, all from Jackson ImmunoResearch. For microtubule depolymerization-regrowth experiments, cells were first incubated at 4°C for 30 min, then rinsed in pre-warmed medium (37°C), followed by incubation at room temperature for 1-2 min to allow microtubule regrowth. Thereafter, cells were fixed and stained as described above.

### Live imaging, ultrastructure expansion microscopy and confocal microscopy

HeLa Centrin-1:GFP cells were transfected with control or TRIM37 siRNAs for 48 hours, transferred to 35mm imaging dishes (Ibidi, cat.no 81156), and imaged at 37°C and 5% CO2 in medium supplemented with 25mM HEPES (Thermofisher) and 1% PenStrep (Thermofisher). Combined DIC and GFP-epifluorescence time-lapse microscopy was performed on a motorized Zeiss Axio Observer D1 using a 63x 1.4 NA plan-Apochromat oil immersion objective, equipped with an Andor Zyla 4.2 sCMOS camera, a piezo controlled Z-stage (Ludl Electronic Products), and an LED light source (Lumencor SOLA II). Imaging was conducted every 10 minutes, capturing Z-stacks of optical sections 0.5um apart, covering a total height of 8 um. Ultrastructure expansion microscopy was conducted essential as reported (Gambarotto et al., 2019). For imaging, the sample was mounted on a 25 mm round poly-D-lysine coated precision coverslip. STED imaging was performed on a Leica TCS SP8 STED 3X microscope with a 100x 1.4 NA oil-immersion objective. Secondary antibodies were 1:500 Alexa-488 (A-11039; Thermofisher) Alexa-594 (ab150072; Abcam) and Atto647N (2418; Hypermol). Confocal images were captured on a Leica TCS SP5 with a HCX PL APO Lambda blue 63× 1.4 NA oil objective. All images shown are maximal intensity projections. Image processing was carried out using Image J and Adobe Photoshop (Adobe).

### Correlative light electron microcopy (CLEM)

HeLa and RPE-1 cells expressing Centrin-1:GFP were cultured in glass-bottom Petri dishes (MatTek, Cat. No. P35G-1.5-14-CGRD), with an alpha-numeric grid pattern, and transfected with control or TRIM37 siRNAs. Cells were chemically fixed 72 h after transfection with a buffered solution of 1 % glutaraldehyde 2 % paraformaldehyde in 0.1 M phosphate buffer at pH 7.4. Dishes were then screened with a wide-field fluorescent microscope (Zeiss Observer D1, using a 63x 1.4 NA oil objective) to identify cells of interest, which were imaged with both transmitted and fluorescence microscopy to register the position of each cell on the grid, as well as the location of their GFP foci, capturing optical slices 500 nm apart. The cells were then washed thoroughly with cacodylate buffer (0.1M, pH 7.4), postfixed for 40 min in 1.0 % osmium tetroxide 1.5% potassium ferrocyanide, and then for 40 min in 1.0% osmium tetroxide alone. Finally, cells were stained for 40 min in 1% uranyl acetate in water before dehydration through increasing concentrations of alcohol and then embedding in Durcupan ACM resin (Fluka, Switzerland). The coverslips were then covered with 1 mm of resin, which was hardened for 18 hours in a 65° C oven. The coverslips were removed from the cured resin by immersing them alternately into hot (60° C) water followed by liquid nitrogen until the coverslips parted. Regions of resin containing the cells of interest were then identified according to their position on the alpha-numeric grid, cut away from the rest of the material and glued to blank resin block. Ultra-thin (50 nm thick) serial sections were cut through the entire cell with a diamond knife (Diatome) and ultramicrotome (Leica Microsystems, UC7), and collected onto single slot grids with a pioloform support film. Sections were further contrasted with lead citrate and uranyl acetate and images taken in a transmission electron microscope (FEI Company, Tecnai Spirit) with a digital camera (FEI Company, Eagle). To correlate the light microscopy images with the EM images and identify the exact position of the Centrin-1:GFP foci, fluorescent images were overlaid onto the electron micrographs of the same cell using Photoshop.

### Statistical analysis

Statistical significance was determined with a Student’s t-test using PRISM software (Graphpad Software Inc.). Statistically significance of pair-wise comparisons are indicated in the figure legends with P < 0.05, P < 0.01 or P < 0.001.

## SUPPLEMENTARY FIGUE LEGENDS

**Suppl. Figure 1.**
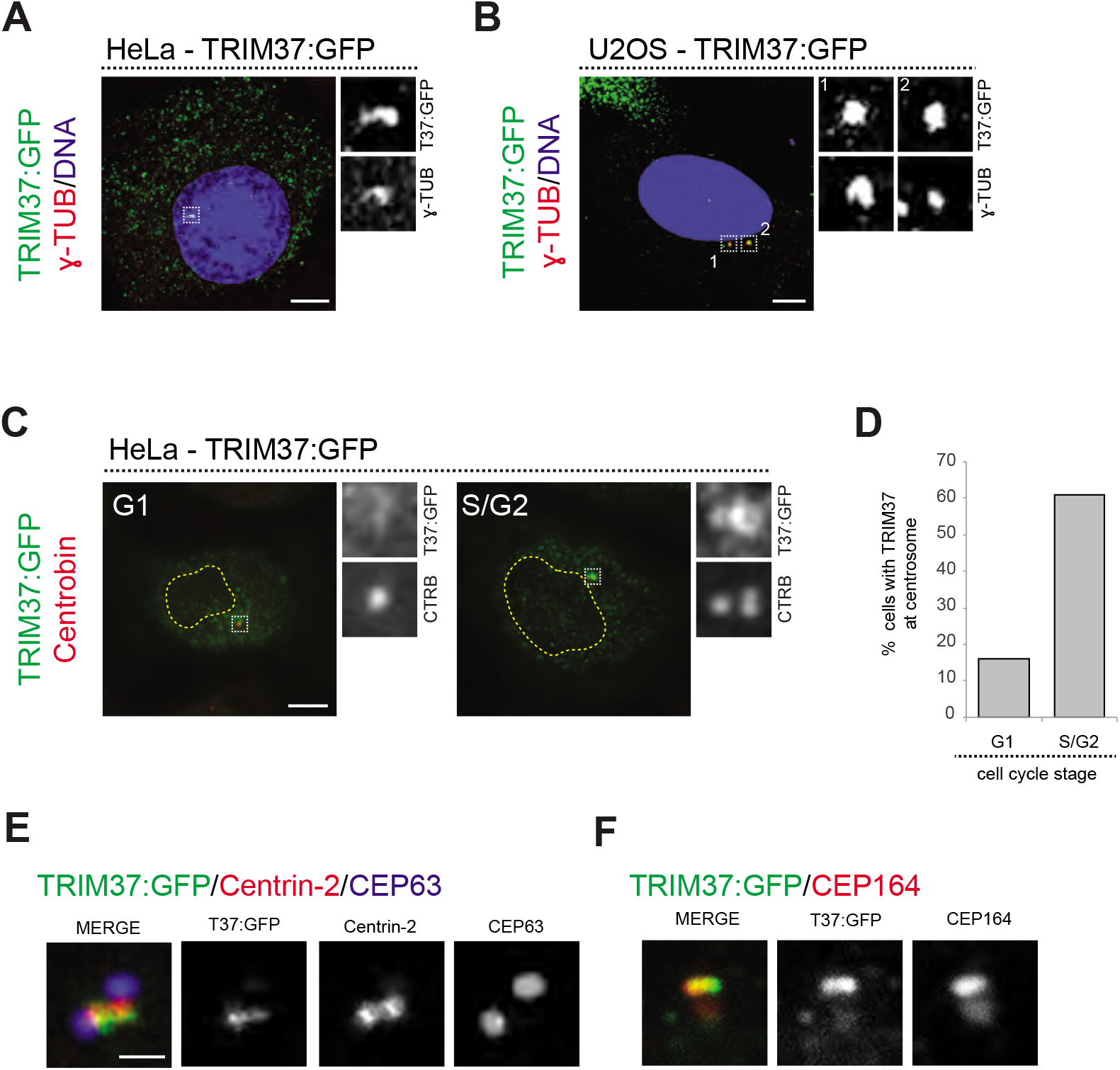
TRIM37:GFP localizes to centrioles. **A**, **B.** HeLa (A) and U2OS (B) cell transfected with TRIM37:GFP, and immunostained for GFP plus γ-tubulin. In this and other supplementary figure panels, scale bars correspond to 5 μm, unless indicated otherwise. **C.** HeLa cells transfected with TRIM37:GFP, and immunostained for GFP plus Centrobin. Cells with a single Centrobin focus (left) were classified as being in G1, cells with two or three Centrobin foci (right) as being in S/G2. **D.** Corresponding percentage of cells in G1 or S/G2 exhibiting TRIM37:GFP at centrosomes. n=50 cells, single experiment. **E, F.** High magnification confocal images of TRIM37:GFP localization with respect to indicated centriolar markers; HeLa cells were fixed 24h after transfection in this case. Scale bar 500 nm.

**Suppl. Figure 2.**
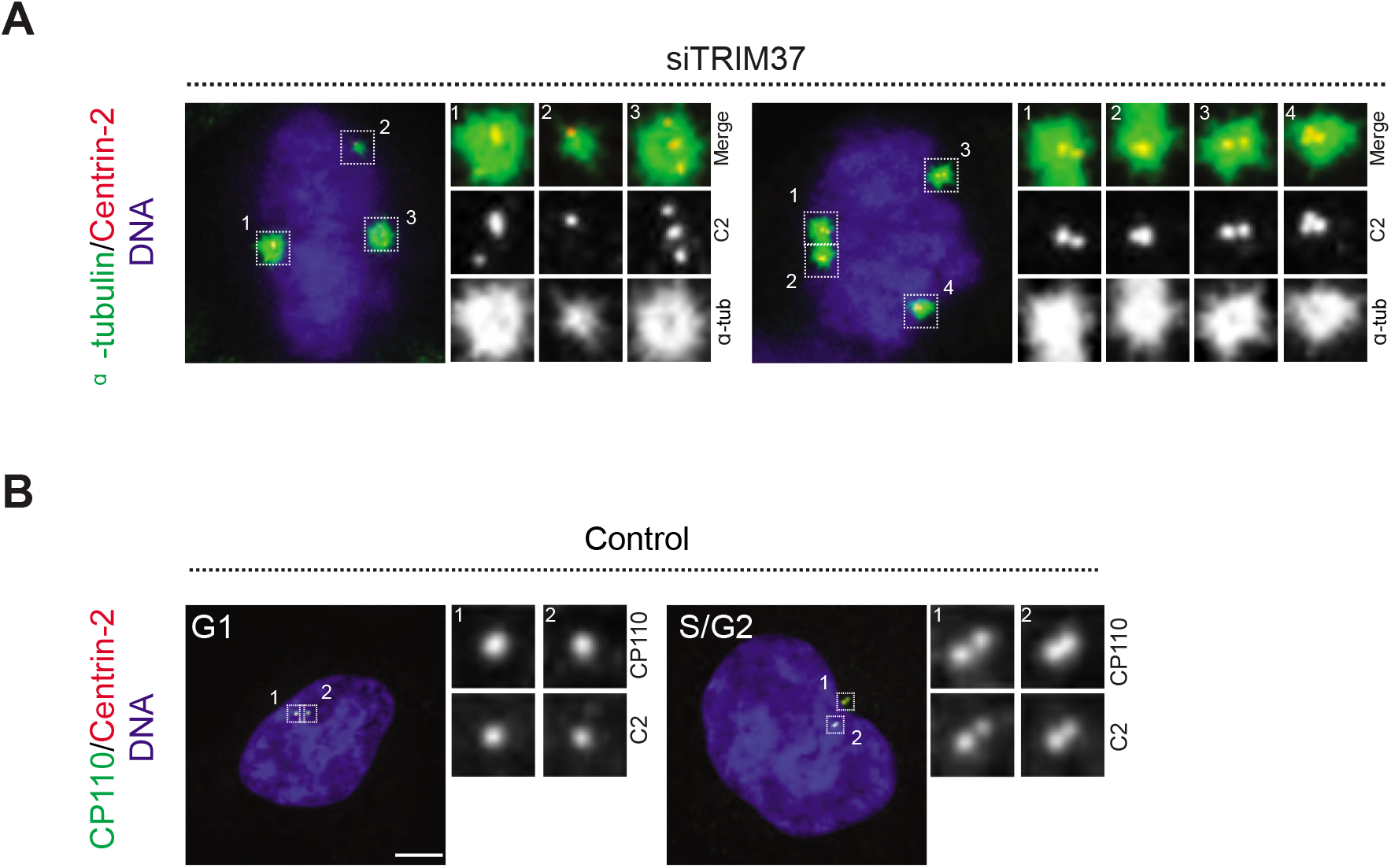
Cenpas can behave as extra MTOCs. **A.** Microtubule depolymerization-regrowth experiment in mitotic HeLa cells treated with control or TRIM37 siRNAs. Microtubules were depolymerized by a 30 min cold shock followed by 1-2 min at room temperature before fixation of cells and immunostaining for Centrin-2 and α-tubulin. **B.** HeLa cells immunostained for Centrin-2 and CP110. Left: G1 cell, with two resident centrioles, right: S/G2 cell with two centriole pairs, each with one resident centriole and one procentriole.

**Supplementary Figure 3.**
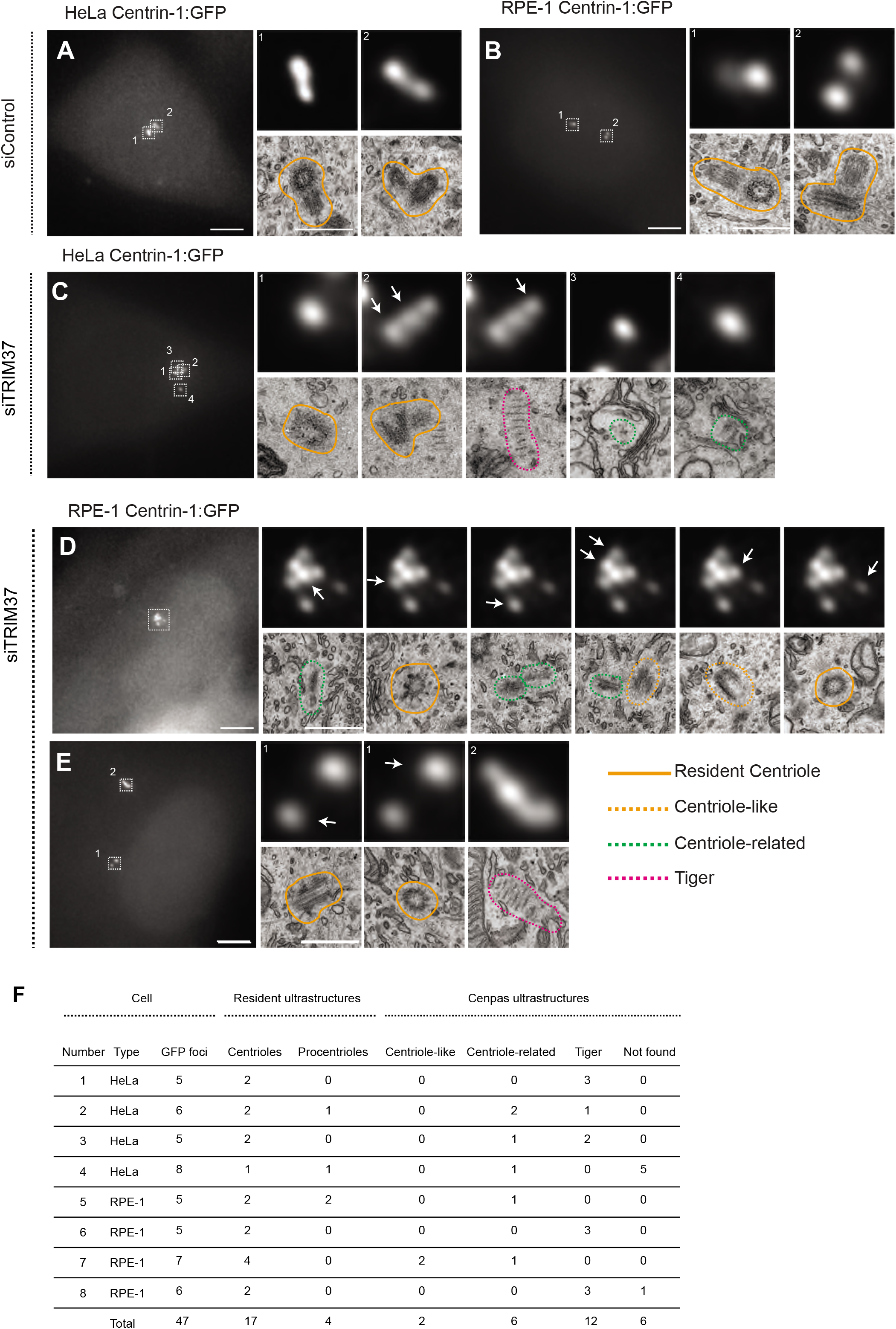
CLEM analysis of Cenpas. **A-E.** CLEM analysis of HeLa and RPE-1 cells expressing Centrin-1:GFP and transfected with control (A-B) or TRIM37 (C-E) siRNAs. Left-most images show maximal intensity projection of wide-field microcopy image covering the entire cell volume; scale bar: 5 μm. Magnified insets from the light microscopy images are shown above the electron microscopy images taken from the corresponding position. C corresponds to cell 2, D to cell 7, and E to cell 6 in Fig. S3F. When present, white arrows indicate the Centrin-1:GFP focus that correlates with the EM imaged shown below. Scale bars in insets are 500 nm. Orange, green and pink dashed lines surround respectively centriole-like, centriole-related and tiger structures. Filled orange lines surround resident centrioles. **F.** Summary of CLEM analysis of HeLa or RPE-1 cells depleted of TRIM37, with number of GFP foci, as well as corresponding resident centriole/procentriole and Cenpas ultrastructure identified by CLEM. See main text for further details. Note that no distinct ultrastructure was found for 5 Centrin-1:GFP foci in cell 4, perhaps reflecting a technical issue in this case.

**Supplementary Figure 4.**
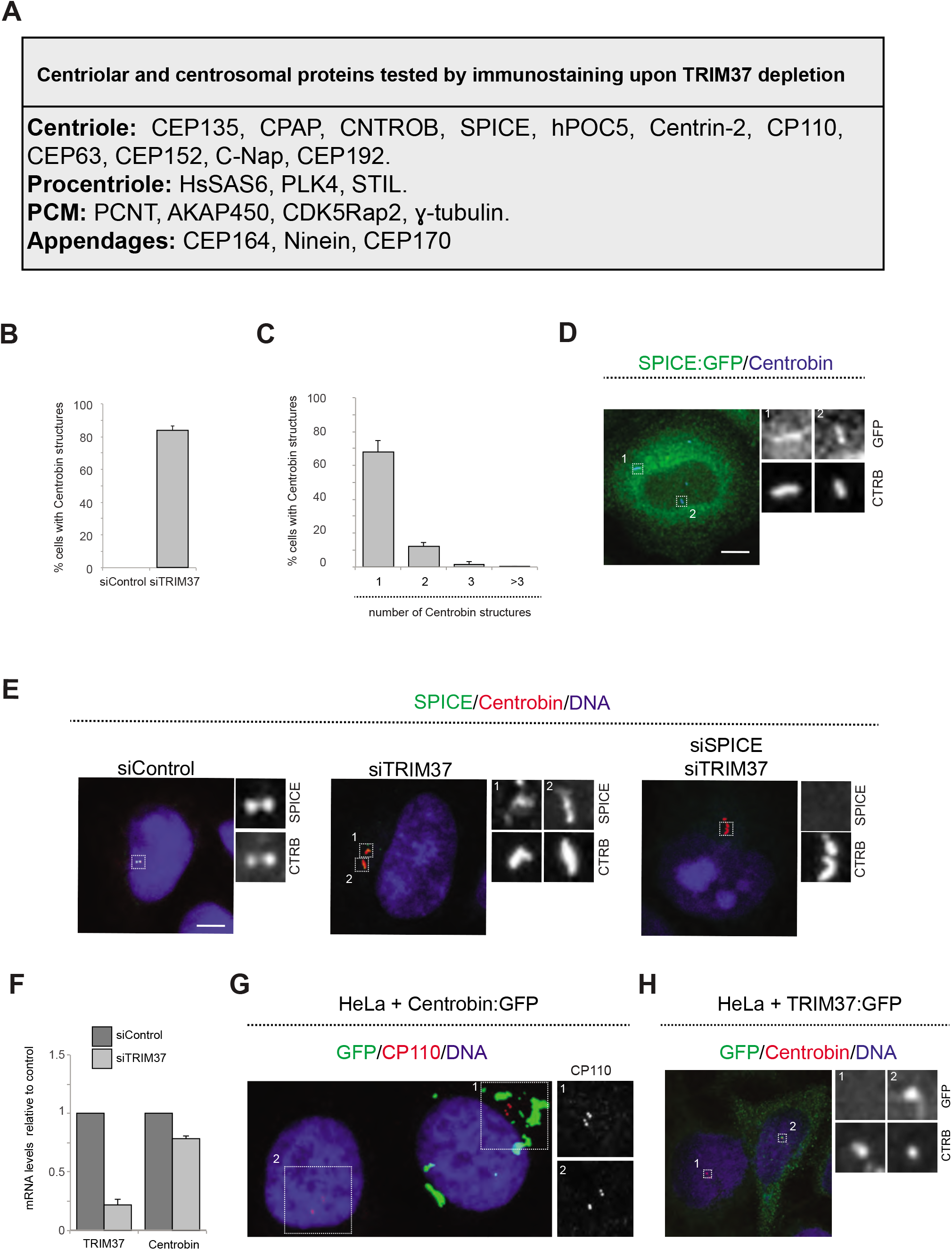
Analysis of Centrobin structures formed upon TRIM37 depletion. **A**, Centriolar and centrosomal proteins analyzed by immunofluorescence upon TRIM37 depletion. See Materials and Methods for antibodies utilized. **B**, **C.** Quantification of frequency (B) and number per cell (C) of Centrobin assemblies in HeLa cells depleted of TRIM37. Unless otherwise indicated, in this and subsequent supplementary figures all graphs report averages from two or more independent experiments (n = 50 cells each), along with SDs. **D.** HeLa cell expressing SPICE:GFP immunostained for GFP and Centrobin. **E.** HeLa cells treated with control, TRIM37 or double TRIM37 and SPICE siRNAs, immunostained for SPICE and Centrobin. **F.** Quantitative real time PCR of TRIM37 and Centrobin mRNA in HeLa cells treated with control or TRIM37 siRNAs. Average of three independent experiments. **G.** HeLa cell (left) and HeLa cell overexpressing Centrobin:GFP (right), immunostained for GFP and CP110. **H.** Confocal images of HeLa cells overexpressing TRIM37:GFP immunostained with antibodies against GFP and Centrobin.

**Supplementary Figure 5.**
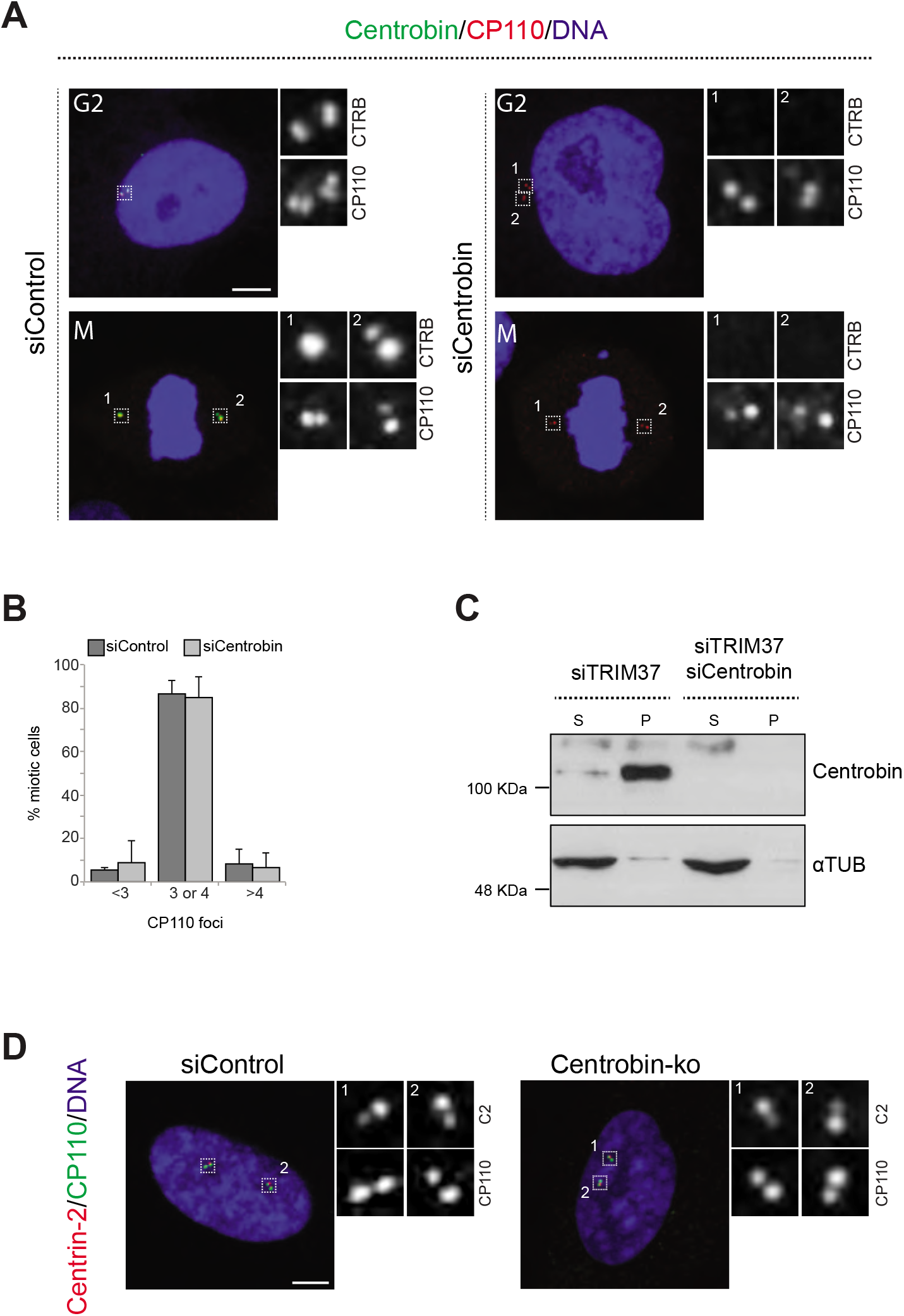
Centrobin is not required for canonical centriole duplication. **A.** HeLa cells in G2 or mitosis, as indicated, treated with control or Centrobin siRNAs, aand immunostained for Centrobin plus CP110. **B.** Corresponding percentage of mitotic cells with indicated number of CP110 foci. **C.** Western blot of soluble (S) and insoluble (P, for pellet) fractions of lysates from HeLa cells transfected with siRNAs against TRIM37 or against both TRIM37 and Centrobin, probed with antibodies against Centrobin (top) or α-tubulin as loading control (bottom). **D.** Control or Centrobin-ko RPE-1 cells transfected with control siRNAs and immunostained for Centrin-2 plus CP110.

**Supplementary Figure 6.**
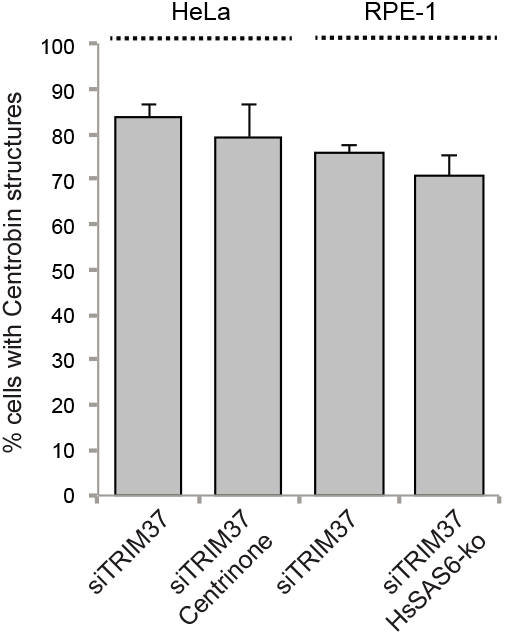
Two pathways contribute to Cenpas formation upon TRIM37 depletion. Percentages of cells with Centrobin structures in the indicated conditions. P value: not significant for both pair-wise comparisons (i.e. siTRIM37 versus Centrinone + siTRIM37 and siTRIM37 versus. HsSAS-6-ko + siTRIM37).

## BIBLIOGRAPHY

Andersen, J.S., Wilkinson, C.J., Mayor, T., Mortensen, P., Nigg, E.A., and Mann, M. (2003). Proteomic characterization of the human centrosome by protein correlation profiling. Nature 426, 570–574.

Archinti, M., Lacasa, C., Teixido-Travesa, N., and Luders, J. (2010). SPICE--a previously uncharacterized protein required for centriole duplication and mitotic chromosome congression. Journal of cell science 123, 3039–3046.

Arquint, C., and Nigg, E.A. (2016). The PLK4-STIL-SAS-6 module at the core of centriole duplication. Biochemical Society transactions 44, 1253–1263.

Avela, K., Lipsanen-Nyman, M., Idanheimo, N., Seemanova, E., Rosengren, S., Makela, T.P., Perheentupa, J., Chapelle, A.D., and Lehesjoki, A.E. (2000). Gene encoding a new RING-B-box-Coiled-coil protein is mutated in mulibrey nanism. Nature genetics 25, 298–301.

Azimzadeh, J., Hergert, P., Delouvee, A., Euteneuer, U., Formstecher, E., Khodjakov, A., and Bornens, M. (2009). hPOC5 is a centrin-binding protein required for assembly of full-length centrioles. The Journal of cell biology 185, 101–114.

Balestra, F.R., Strnad, P., Fluckiger, I., and Gönczy, P. (2013). Discovering regulators of centriole biogenesis through siRNA-based functional genomics in human cells. Developmental cell 25, 555–571.

Banterle, N., and Gönczy, P. (2017). Centriole Biogenesis: From Identifying the Characters to Understanding the Plot. Annual review of cell and developmental biology 33, 23–49.

Bettencourt-Dias, M., Hildebrandt, F., Pellman, D., Woods, G., and Godinho, S.A. (2011). Centrosomes and cilia in human disease. Trends Genet 27, 307–315.

Bhatnagar, S., Gazin, C., Chamberlain, L., Ou, J., Zhu, X., Tushir, J.S., Virbasius, C.M., Lin, L., Zhu, L.J., Wajapeyee, N. et al. (2014). TRIM37 is a new histone H2A ubiquitin ligase and breast cancer oncoprotein. Nature 516, 116–120.

Bornens, M. (2012). The centrosome in cells and organisms. Science 335, 422–426.

Brown, N.J., Marjanovic, M., Luders, J., Stracker, T.H., and Costanzo, V. (2013). Cep63 and cep152 cooperate to ensure centriole duplication. PloS one 8, e69986.

Chavali, P.L., Putz, M., and Gergely, F. (2014). Small organelle, big responsibility: the role of centrosomes in development and disease. Philosophical transactions of the Royal Society of London Series B, Biological sciences 369.

Comartin, D., Gupta, G.D., Fussner, E., Coyaud, E., Hasegan, M., Archinti, M., Cheung, S.W., Pinchev, D., Lawo, S., Raught, B., et al. (2013). CEP120 and SPICE1 cooperate with CPAP in centriole elongation. Current biology : CB 23, 1360–1366.

Courtois, A., Schuh, M., Ellenberg, J., and Hiiragi, T. (2012). The transition from meiotic to mitotic spindle assembly is gradual during early mammalian development. The Journal of cell biology 198, 357–370.

Duensing, A., Liu, Y., Perdreau, S.A., Kleylein-Sohn, J., Nigg, E.A., and Duensing, S. (2007). Centriole overduplication through the concurrent formation of multiple daughter centrioles at single maternal templates. Oncogene 26, 6280–6288.

Fritz-Laylin, L.K., Levy, Y.Y., Levitan, E., Chen, S., Cande, W.Z., Lai, E.Y., and Fulton, C. (2016). Rapid centriole assembly in Naegleria reveals conserved roles for both de novo and mentored assembly. Cytoskeleton 73, 109–116.

Fulton, C., and Dingle, A.D. (1971). Basal bodies, but not centrioles, in Naegleria. The Journal of cell biology 51, 826–836.

Gambarotto, D., Zwettler, F.U., Le Guennec, M., Schmidt-Cernohorska, M., Fortun, D., Borgers, S., Heine, J., Schloetel, J.G., Reuss, M., Unser, M., et al. (2019). Imaging cellular ultrastructures using expansion microscopy (U-ExM). Nature methods 16, 71–74.

Ganem, N.J., Godinho, S.A., and Pellman, D. (2009). A mechanism linking extra centrosomes to chromosomal instability. Nature 460, 278–282.

Godinho, S.A., Picone, R., Burute, M., Dagher, R., Su, Y., Leung, C.T., Polyak, K., Brugge, J.S., Thery, M., and Pellman, D. (2014). Oncogene-like induction of cellular invasion from centrosome amplification. Nature 510, 167–171.

Gönczy, P. (2012). Towards a molecular architecture of centriole assembly. Nature reviews Molecular cell biology 13, 425–435.

Gönczy, P. (2015). Centrosomes and cancer: revisiting a long-standing relationship. Nature reviews Cancer 15, 639–652.

Gönczy, P., and Hatzopoulos, G.N. (2019). Centriole assembly at a glance. Journal of cell science 132.

Gudi, R., Zou, C., Li, J., and Gao, Q. (2011). Centrobin-tubulin interaction is required for centriole elongation and stability. The Journal of cell biology 193, 711–725.

Guichard, P., Hamel, V., and Gonczy, P. (2018). The Rise of the Cartwheel: Seeding the Centriole Organelle. BioEssays : news and reviews in molecular, cellular and developmental biology 40, e1700241.

Guichard, P., Hamel, V., Le Guennec, M., Banterle, N., Iacovache, I., Nemcikova, V., Fluckiger, I., Goldie, K.N., Stahlberg, H., Levy, D., et al. (2017). Cell-free reconstitution reveals centriole cartwheel assembly mechanisms. Nat Commun 8, 14813.

Habedanck, R., Stierhof, Y.D., Wilkinson, C.J., and Nigg, E.A. (2005). The Polo kinase Plk4 functions in centriole duplication. Nature cell biology 7, 1140–1146.

Hirono, M. (2014). Cartwheel assembly. Philosophical transactions of the Royal Society of London Series B, Biological sciences 369.

Hu, C.E., and Gan, J. (2017). TRIM37 promotes epithelialmesenchymal transition in colorectal cancer. Molecular medicine reports 15, 1057–1062.

Jakobsen, L., Vanselow, K., Skogs, M., Toyoda, Y., Lundberg, E., Poser, I., Falkenby, L.G., Bennetzen, M., Westendorf, J., Nigg, E.A., et al. (2011). Novel asymmetrically localizing components of human centrosomes identified by complementary proteomics methods. The EMBO journal 30, 1520–1535.

Jeong, Y., Lee, J., Kim, K., Yoo, J.C., and Rhee, K. (2007). Characterization of NIP2/centrobin, a novel substrate of Nek2, and its potential role in microtubule stabilization. Journal of cell science 120, 2106–2116.

Jiang, J., Yu, C., Chen, M., Tian, S., and Sun, C. (2015). Over-expression of TRIM37 promotes cell migration and metastasis in hepatocellular carcinoma by activating Wnt/beta-catenin signaling. Biochemical and biophysical research communications 464, 1120–1127.

Kallijarvi, J., Avela, K., Lipsanen-Nyman, M., Ulmanen, I., and Lehesjoki, A.E. (2002). The TRIM37 gene encodes a peroxisomal RING-B-box-coiled-coil protein: classification of mulibrey nanism as a new peroxisomal disorder. American journal of human genetics 70, 1215–1228.

Kallijarvi, J., Lahtinen, U., Hamalainen, R., Lipsanen-Nyman, M., Palvimo, J.J., and Lehesjoki, A.E. (2005). TRIM37 defective in mulibrey nanism is a novel RING finger ubiquitin E3 ligase. Exp Cell Res 308, 146–155.

Karlberg, N., Karlberg, S., Karikoski, R., Mikkola, S., Lipsanen-Nyman, M., and Jalanko, H. (2009). High frequency of tumours in Mulibrey nanism. The Journal of pathology 218, 163–171.

Kettunen, K.M., Karikoski, R., Hamalainen, R.H., Toivonen, T.T., Antonenkov, V.D., Kulesskaya, N., Voikar, V., Holtta-Vuori, M., Ikonen, E., Sainio, K., et al. (2016). Trim37-deficient mice recapitulate several features of the multi-organ disorder Mulibrey nanism. Biology open.

Khodjakov, A., Rieder, C.L., Sluder, G., Cassels, G., Sibon, O., and Wang, C.L. (2002). De novo formation of centrosomes in vertebrate cells arrested during S phase. The Journal of cell biology 158, 1171–1181.

Kitagawa, D., Kohlmaier, G., Keller, D., Strnad, P., Balestra, F.R., Fluckiger, I., and Gönczy, P. (2011a). Spindle positioning in human cells relies on proper centriole formation and on the microcephaly proteins CPAP and STIL. Journal of cell science 124, 3884–3893.

Kitagawa, D., Vakonakis, I., Olieric, N., Hilbert, M., Keller, D., Olieric, V., Bortfeld, M., Erat, M.C., Fluckiger, I., Gönczy, P., et al. (2011b). Structural basis of the 9-fold symmetry of centrioles. Cell 144, 364–375.

Klebba, J.E., Buster, D.W., McLamarrah, T.A., Rusan, N.M., and Rogers, G.C. (2015). Autoinhibition and relief mechanism for Polo-like kinase 4. Proceedings of the National Academy of Sciences of the United States of America 112, E657–666.

Kohlmaier, G., Loncarek, J., Meng, X., McEwen, B.F., Mogensen, M.M., Spektor, A., Dynlacht, B.D., Khodjakov, A., and Gönczy, P. (2009). Overly long centrioles and defective cell division upon excess of the SAS-4-related protein CPAP. Current biology : CB 19, 1012–1018.

La Terra, S., English, C.N., Hergert, P., McEwen, B.F., Sluder, G., and Khodjakov, A. (2005). The de novo centriole assembly pathway in HeLa cells: cell cycle progression and centriole assembly/maturation. The Journal of cell biology 168, 713–722.

Levine, M.S., Bakker, B., Boeckx, B., Moyett, J., Lu, J., Vitre, B., Spierings, D.C., Lansdorp, P.M., Cleveland, D.W., Lambrechts, D., et al. (2017). Centrosome Amplification Is Sufficient to Promote Spontaneous Tumorigenesis in Mammals. Developmental cell 40, 313–322 e315.

Li, J., Kim, S., Kobayashi, T., Liang, F.X., Korzeniewski, N., Duensing, S., and Dynlacht, B.D. (2012). Neurl4, a novel daughter centriole protein, prevents formation of ectopic microtubule organizing centres. EMBO reports 13, 547–553.

Loncarek, J., Hergert, P., and Khodjakov, A. (2010). Centriole reduplication during prolonged interphase requires procentriole maturation governed by Plk1. Current biology : CB 20, 1277–1282.

Lukinavicius, G., Lavogina, D., Orpinell, M., Umezawa, K., Reymond, L., Garin, N., Gonczy, P., and Johnsson, K. (2013). Selective chemical crosslinking reveals a Cep57-Cep63-Cep152 centrosomal complex. Current biology : CB 23, 265–270.

Mahjoub, M.R., and Stearns, T. (2012). Supernumerary centrosomes nucleate extra cilia and compromise primary cilium signaling. Current biology : CB 22, 1628–1634.

Meitinger, F., Anzola, J.V., Kaulich, M., Richardson, A., Stender, J.D., Benner, C., Glass, C.K., Dowdy, S.F., Desai, A., Shiau, A.K., et al. (2016). 53BP1 and USP28 mediate p53 activation and G1 arrest after centrosome loss or extended mitotic duration. The Journal of cell biology 214, 155–166.

Montenegro Gouveia, S., Zitouni, S., Kong, D., Duarte, P., Ferreira Gomes, B., Sousa, A.L., Tranfield, E.M., Hyman, A., Loncarek, J., and Bettencourt-Dias, M. (2018). PLK4 is a microtubule-associated protein that self-assembles promoting de novo MTOC formation. Journal of cell science 132.

Moyer, T.C., Clutario, K.M., Lambrus, B.G., Daggubati, V., and Holland, A.J. (2015). Binding of STIL to Plk4 activates kinase activity to promote centriole assembly. The Journal of cell biology 209, 863–878.

Moyer, T.C., and Holland, A.J. (2019). PLK4 promotes centriole duplication by phosphorylating STIL to link the procentriole cartwheel to the microtubule wall. eLife 8.

Nigg, E.A., and Holland, A.J. (2018). Once and only once: mechanisms of centriole duplication and their deregulation in disease. Nature reviews Molecular cell biology.

Nigg, E.A., and Raff, J.W. (2009). Centrioles, centrosomes, and cilia in health and disease. Cell 139, 663–678.

Ogungbenro, Y.A., Tena, T.C., Gaboriau, D., Lalor, P., Dockery, P., Philipp, M., and Morrison, C.G. (2018). Centrobin controls primary ciliogenesis in vertebrates. The Journal of cell biology 217, 1205–1215.

Ohta, M., Ashikawa, T., Nozaki, Y., Kozuka-Hata, H., Goto, H., Inagaki, M., Oyama, M., and Kitagawa, D. (2014). Direct interaction of Plk4 with STIL ensures formation of a single procentriole per parental centriole. Nat Commun 5, 5267.

Piel, M., Meyer, P., Khodjakov, A., Rieder, C.L., and Bornens, M. (2000). The respective contributions of the mother and daughter centrioles to centrosome activity and behavior in vertebrate cells. The Journal of cell biology 149, 317–330.

Schmidt, T.I., Kleylein-Sohn, J., Westendorf, J., Le Clech, M., Lavoie, S.B., Stierhof, Y.D., and Nigg, E.A. (2009). Control of centriole length by CPAP and CP110. Current biology : CB 19, 1005–1011.

Sercin, O., Larsimont, J.C., Karambelas, A.E., Marthiens, V., Moers, V., Boeckx, B., Le Mercier, M., Lambrechts, D., Basto, R., and Blanpain, C. (2016). Transient PLK4 overexpression accelerates tumorigenesis in p53-deficient epidermis. Nature cell biology 18, 100–110.

Shin, W., Yu, N.K., Kaang, B.K., and Rhee, K. (2015). The microtubule nucleation activity of centrobin in both the centrosome and cytoplasm. Cell cycle 14, 1925–1931.

Shiratsuchi, G., Takaoka, K., Ashikawa, T., Hamada, H., and Kitagawa, D. (2015). RBM14 prevents assembly of centriolar protein complexes and maintains mitotic spindle integrity. The EMBO journal 34, 97–114.

Sillibourne, J.E., Tack, F., Vloemans, N., Boeckx, A., Thambirajah, S., Bonnet, P., Ramaekers, F.C., Bornens, M., and Grand-Perret, T. (2010). Autophosphorylation of polo-like kinase 4 and its role in centriole duplication. Molecular biology of the cell 21, 547–561.

Sinclair, C.S., Rowley, M., Naderi, A., and Couch, F.J. (2003). The 17q23 amplicon and breast cancer. Breast cancer research and treatment 78, 313–322.

Strnad, P., Leidel, S., Vinogradova, T., Euteneuer, U., Khodjakov, A., and Gönczy, P. (2007). Regulated HsSAS-6 levels ensure formation of a single procentriole per centriole during the centrosome duplication cycle. Developmental cell 13, 203–213.

Sullenberger, C., Vasquez-Limeta, A., Kong, D., and Loncarek, J. (2020). With Age Comes Maturity: Biochemical and Structural Transformation of a Human Centriole in the Making. Cells 9.

Tang, C.J., Fu, R.H., Wu, K.S., Hsu, W.B., and Tang, T.K. (2009). CPAP is a cell-cycle regulated protein that controls centriole length. Nature cell biology 11, 825–831.

Thauvin-Robinet, C., Lee, J.S., Lopez, E., Herranz-Perez, V., Shida, T., Franco, B., Jego, L., Ye, F., Pasquier, L., Loget, P., et al. (2014). The oral-facial-digital syndrome gene C2CD3 encodes a positive regulator of centriole elongation. Nature genetics 46, 905–911.

Tsou, M.F., Wang, W.J., George, K.A., Uryu, K., Stearns, T., and Jallepalli, P.V. (2009). Polo kinase and separase regulate the mitotic licensing of centriole duplication in human cells. Developmental cell 17, 344–354.

van Breugel, M., Hirono, M., Andreeva, A., Yanagisawa, H.A., Yamaguchi, S., Nakazawa, Y., Morgner, N., Petrovich, M., Ebong, I.O., Robinson, C.V., et al. (2011). Structures of SAS-6 suggest its organization in centrioles. Science 331, 1196–1199.

Wang, W., Xia, Z.J., Farre, J.C., and Subramani, S. (2017). TRIM37, a novel E3 ligase for PEX5-mediated peroxisomal matrix protein import. The Journal of cell biology 216, 2843–2858.

Wang, W.J., Acehan, D., Kao, C.H., Jane, W.N., Uryu, K., and Tsou, M.F. (2015). De novo centriole formation in human cells is error-prone and does not require SAS-6 self-assembly. eLife 4.

Wen, W., Meinkoth, J.L., Tsien, R.Y., and Taylor, S.S. (1995). Identification of a signal for rapid export of proteins from the nucleus. Cell 82, 463–473.

Wong, Y.L., Anzola, J.V., Davis, R.L., Yoon, M., Motamedi, A., Kroll, A., Seo, C.P., Hsia, J.E., Kim, S.K., Mitchell, J.W., et al. (2015). Cell biology. Reversible centriole depletion with an inhibitor of Polo-like kinase 4. Science 348, 1155–1160.

Zou, C., Li, J., Bai, Y., Gunning, W.T., Wazer, D.E., Band, V., and Gao, Q. (2005). Centrobin: a novel daughter centriole-associated protein that is required for centriole duplication. The Journal of cell biology 171, 437–445.

